# Functional and pangenomic exploration of Roc two-component regulatory systems identifies novel players across *Pseudomonas* species

**DOI:** 10.1101/2024.10.18.618891

**Authors:** Victor Simon, Julian Trouillon, Ina Attrée, Sylvie Elsen

## Abstract

The opportunistic pathogen *Pseudomonas aeruginosa* counts on a large collection of two-component regulatory systems (TCSs) to sense and adapt to changing environments. Among them, the Roc (<u>R</u>egulation <u>o</u>f *<u>c</u>up*) system is a one-of-a-kind network of branched TCSs, composed of two histidine kinases (HKs) (RocS1 and RocS2) interacting with three response regulators (RRs) (RocA1, RocR and RocA2), which regulate virulence, antibiotic resistance and biofilm formation. Based on extensive work on the Roc system, previous data suggested the existence of other key regulators yet to be discovered. In this work, we identified PA4080, renamed RocA3, as a fourth RR that is activated by RocS1 and RocS2 and that positively controls the expression of the *cupB* operon. Comparative genomic analysis of the locus identified a gene - *rocR3* - adjacent to *rocA3* in a subpopulation of strains which encodes a protein with structural and functional similarity to the c-di-GMP phosphodiesterase RocR. Furthermore, we identified a fourth branch of the Roc system consisting of the PA2583 HK, renamed RocS4, and of the Hpt protein HptA. Using a bacterial two-hybrid system, we showed that RocS4 interacts with HptA, which in turn interacts with RocA1, RocA2 and RocR3. Finally, we mapped the pangenomic RRs repertoire establishing a comprehensive view of the plasticity of such regulators among clades of the species. Overall, our work provides a comprehensive inter-species definition of the Roc system, nearly doubling the number of proteins known to be involved in this interconnected network of TCSs controlling pathogenicity in *Pseudomonas* species.

**GRAPHICAL ABSTRACT:** 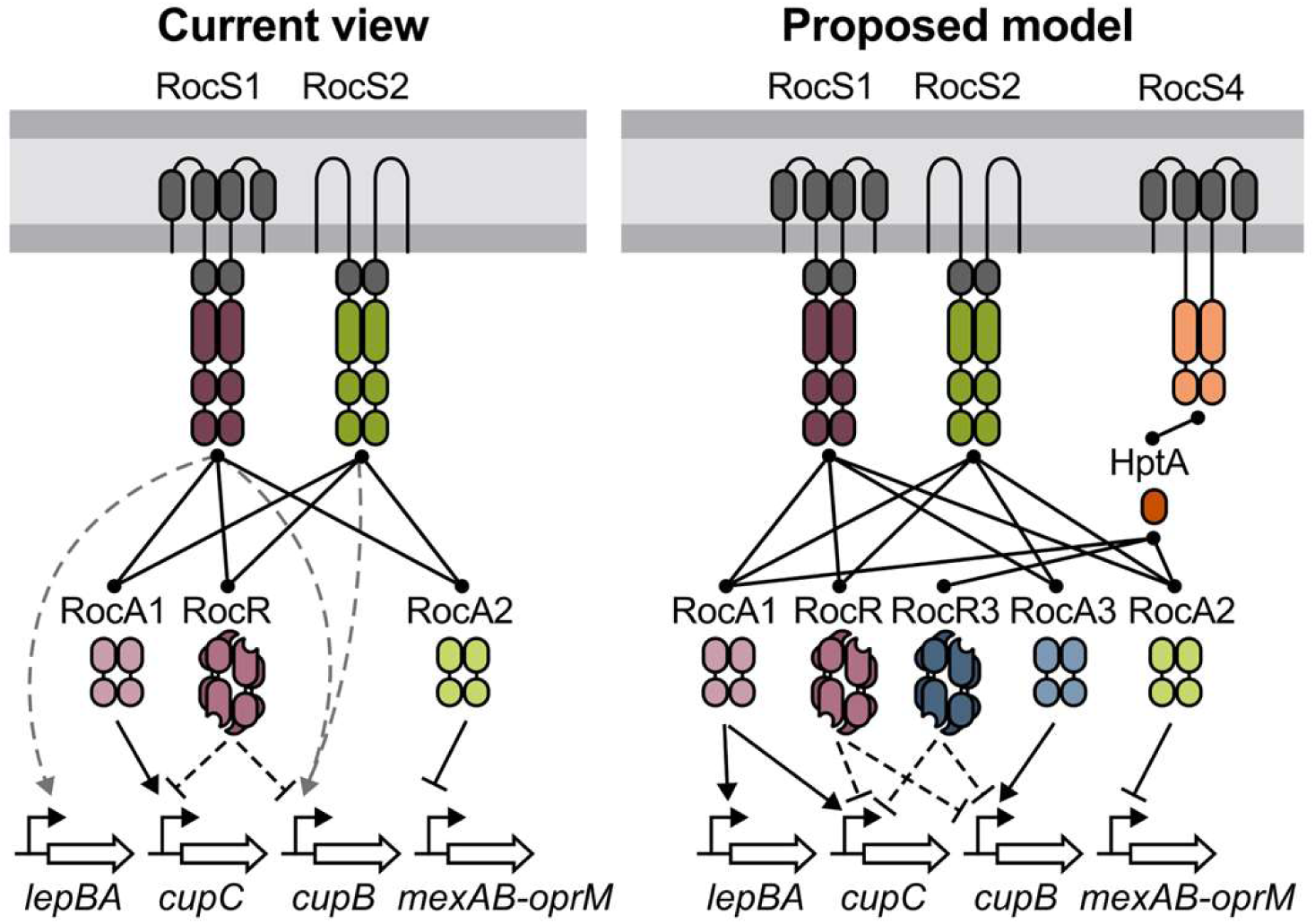

**ABBREVIATED SUMMARY:** Roc system account for a particularly interconnected yet incomplete network of two-component regulatory system involved in the virulence of *Pseudomonas aeruginosa*. Our work identified the missing RocA3 regulator and propose new players of the system delineating their conservation between the clade of the species.

## Introduction

*Pseudomonas aeruginosa* is an opportunistic pathogen responsible for a wide range of human infections, whose variability in virulence and antibiotic resistance capacity relies mainly on the high heterogeneity of its large genomes (Klockgether and Tummler, 2017). Phylogenetic analyses determined the population structure of the species, identifying 5 different clades with, in particular, distinct pathogenesis strategies (Freschi *et al*., 2019; Ozer *et al*., 2019; Quiroz-Morales *et al*., 2022). Clade 3 includes the PA7-like strains known as taxonomic outliers (Roy *et al*., 2010), which have been recently reclassified as a new species, *P. paraeruginosa* (Rudra *et al*., 2022). The ability of these species to cause disease and survive in diverse environments is largely due to sophisticated transcriptional regulatory networks, of which two-component regulatory systems (TCSs) are key players (Galán-Vásquez *et al*., 2011; Francis *et al*., 2017).

TCSs are widespread signaling machines that allow bacteria to directly respond to environmental cues and adapt to diverse conditions (Stock *et al*., 2000; Capra and Laub, 2012; Zschiedrich *et al*., 2016). As their name suggests, classical TCSs are a pair of signal transduction proteins communicating by phosphotransfer: a sensor histidine kinase (HK), commonly transmembrane, and a cytoplasmic response regulator (RR). HKs are often bifunctional enzymes with phosphatase and autokinase activities that are modulated by one or more chemical or physical signals, most of which are still unknown (Zschiedrich *et al*., 2016). Upon signal detection through its input domain(s), the HK usually autophosphorylates a conserved histidine residue within its transmitter domain (H domain). The phosphate is then transferred to a conserved aspartate residue on the receiver domain (D or REC domain) of the RR, resulting in its activation (Stock *et al*., 2000). Since most RRs are transcription factors, the output response is often the modification of target gene expression.

Besides the typical TCSs with a two-step phosphate transfer, there are many reports of four-step phosphorelays where the phosphate is transferred between additional domains on the HK or other protein partners (Stock *et al*., 2000; Zschiedrich *et al*., 2016). For instance, an “unorthodox” HK has two additional C-terminal domains, a receiver domain and a histidine-containing phosphotransfer (Hpt) domain in a so-called H1-D1-H2 organization. In this case, internal phosphate transfer along the H1-D1-H2 domains precedes phosphorylation of the corresponding RR, as exemplified by GacS in *P. aeruginosa*, which plays a crucial role in the transition from chronic to acute infection (Goodman *et al*., 2004). Four-step phosphorelays also occur in the case of a “hybrid” HK that contains an additional D1 receiver domain and requires an external Hpt protein to phosphorylate the RR. In some cases, multiple interacting partners are involved, where a RR can be activated by several HKs or where one HK can activate several RRs, further increasing the complexity of these systems (Francis and Porter, 2019). Gene duplication, with subsequent divergence, and acquisition by horizontal gene transfer (HGT) are the main sources of new pathways (Alm *et al*., 2006; Capra and Laub, 2012). *In silico* modeling suggests that rapid isolation of signaling between newly duplicated TCSs is necessary to limit non-cognate HK and RR interactions (i. e. cross-talk) that are a potential burden on cell fitness (Capra and Laub, 2012; Rowland and Deeds, 2014).

Approximately 130 genes that code for TCS partners have been predicted in the genome of the *P. aeruginosa* strain PAO1 (Rodrigue *et al*., 2000), which emerged through both recruitment and gene duplication events (Chen *et al*., 2004). Many of these TCSs have been shown to be involved in key infection-related processes, including motility, biofilm formation, cytotoxicity, virulence and antibiotic resistance (Francis *et al*., 2017; Sultan *et al*., 2021). Among them, the Roc (<u>r</u>egulation <u>o</u>f *<u>c</u>up*) signaling pathway in *P. aeruginosa* stands out for its large number of interconnected players. It consists of two paralogous unorthodox HKs (RocS1 and RocS2) interacting with three RRs (RocA1, RocA2 and RocR) (Kulasekara *et al*., 2004; Kuchma *et al*., 2005; Sivaneson *et al*., 2011). The proteins are encoded by two distinct genetic loci, the Roc1 system by *rocA1* and the divergently transcribed *rocR-rocS1* operon, and the Roc2 system by the *rocA2-rocS2* operon. Despite these differences in genetic organization, phylogenetic studies indicated that the two loci evolved by gene duplication and probable gene rearrangement (Chen *et al*., 2004). This common origin may underlie why both HKs RocS1 and RocS2 can interact with and activate all three Roc RRs (Kulasekara *et al*., 2004; Sivaneson *et al*., 2011). Of these three RRs, RocA1 and RocA2 are transcription factors and RocR is a c-di-GMP phosphodiesterase (Rao *et al*., 2004; Kulasekara *et al*., 2004; Sivaneson *et al*., 2011). The main targets of the Roc system are two operons encoding Cup (<u>c</u>haperone-<u>u</u>sher <u>p</u>athway) fimbriae, surface appendages that participate in biofilm formation through adhesion to abiotic surfaces, cell-cell interaction and microcolony formation (Ruer *et al*., 2008; Giraud *et al*., 2009; Giraud *et al*., 2011). Both the *cupC* and *cupB* operons were shown to be activated in a RocS1- and RocS2-dependent manner. However, whereas RocA1 controls *cupC* expression, none of the three known Roc RRs accounts for *cupB* activation: this suggests the existence of an unknown additional regulator of the Roc system (Kulasekara *et al*., 2004; Sivaneson *et al*., 2011).

Recently, we explored the global TCSs regulatory network in different *Pseudomonas* strains by characterizing *in vitro* DNA-binding sites for 55 RRs using DAP-seq (DNA affinity purification and sequencing) (Trouillon *et al*., 2021). The study was carried out on three reference strains, *P. aeruginosa* PAO1 and PA14 (Freschi *et al*., 2019), and *P. paraeruginosa* IHMA87 (Kos *et al*., 2015), allowing the intra- and inter-species comparison of regulatory networks through the 48 regulators that are conserved within the three strains (Trouillon *et al*., 2021). In the present study, we re-examined the DAP-seq data to find the RRs that bind to the promoter of the *cupB* operon and found that the uncharacterized orphan RR PA4080 exhibited one of the strongest bindings on the PA14 genome (Trouillon *et al*., 2021). We experimentally demonstrated that PA4080, whose gene is located downstream of the *cupB* operon, activates this operon in a RocS1- and RocS2-dependent manner, and was therefore named RocA3. Using comparative genomics, we identified several new putative partners, either conserved or restricted to a few clades of *P. aeruginosa* and *P. paraeruginosa*. Our work expands our understanding of the Roc system and highlights how such interconnected TCSs contribute to the diversity of regulatory networks in these species.

## Results

### PA4080 is an activator of the *cupB* operon

Previous DAP-seq data (Trouillon *et al*., 2021) were re-analysed to find potential RRs that control the expression of the *cupB* operon. One of the strongest peak enrichment upstream of the *cupB1* gene, the first gene of the *cupB* operon, was observed for RR PA4080 (**Fig. 1A**), with the binding signal centered at 214 bp from the coding sequence (**Fig. S1A**) (Trouillon *et al*., 2021). This binding was only observed on the PA14 genome as DAP-seq with PA4080 did not provide enriched targets on the PAO1 and IHMA87 genomes, which may be due to a technical problem with the samples. The *PA4080* gene, which is conserved across *P. aeruginosa* and *P. paraeruginosa* strains, is encoded downstream of the *cupB* operon, and this physical proximity strengthens a hypothetical regulatory link (Lawrence, 2003). To first assess the regulatory role of PA4080 on the expression of the *cupB* operon, a transcriptional *lacZ* fusion with the *cupB1* promoter (P*cupB1*-*lacZ*) was constructed and integrated into the PAO1 chromosome, while *PA4080* was overexpressed through the arabinose-inducible P*BAD* promoter on the pJN105 replicative plasmid. As previously observed for the different *cup* operons (Kulasekara *et al*., 2004), the *cupB* operon is poorly expressed under laboratory conditions, with a level of β-galactosidase activity of the transcriptional fusion below 10 Miller units (**Fig. 1B**). However, overexpression of *PA4080* strongly induced *cupB1* expression with a 350-fold increase in reporter activity, demonstrating that PA4080 is an activator of this operon. Since the RR was also able to bind upstream of its own gene *in vitro* (**Fig. 1A**), we used a similar approach and observed that PA4080 positively autoregulates its own gene (**Fig. 1B**). Two known targets of the Roc system controlled by either RocA1 or RocA2 were also tested by generating and analyzing the expression of P*cupC1*-*lacZ* (activated by RocA1; Kulasekara *et al*., 2004) and P*mexA*-*lacZ* (inhibited by RocA2; Sivaneson *et al*., 2011). Our data showed that PA4080 does not control the expression of the *cupC* and *mexAB*-*oprM* operons (**Fig. S1B**).

**FIG 1.**
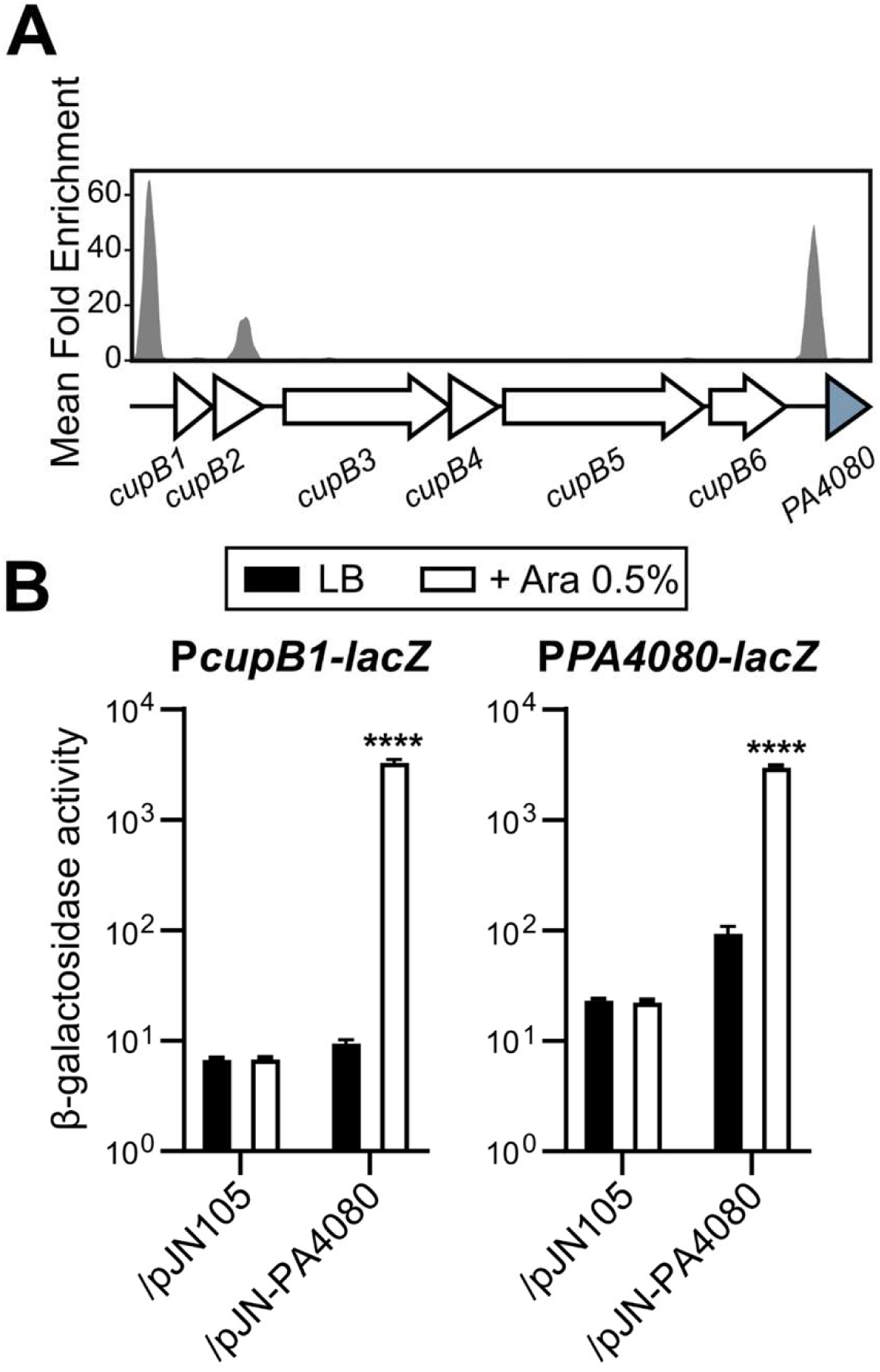
PA4080 regulates the *cupB* operon and its own expression. (A) Reanalyses of previously published data on the PA14 genome (Trouillon *et al*., 2021) showing the enrichment coverage track of DAP-seq against the negative control for the RR PA4080 (PA14_11120) at the indicated locus. (B) β-galactosidase activities of the indicated strains harboring the P*cupB1-lacZ* and P*PA4080-lacZ* transcriptional fusions. The strains also carried either the empty pJN105 or the pJN-PA4080 plasmid, and expression of *PA4080* was induced with 0.5% arabinose for 2.5 h in LB medium. Experiments were performed in triplicate and the error bars represent the SEM. Statistical analysis was performed using two-way ANOVA, followed by Dunnett’s test for comparison with the control condition (PAO1 WT /pJN105 in LB). \*\*\*\**p* < 0.0001.

Taken together, our results demonstrated that PA4080 is a direct activator of the *cupB* operon and its own gene. Considering the incomplete regulatory system already known to activate *cupB* expression, we hypothesized that PA4080 is the suspected additional regulator of the Roc system (**Fig. 2A**).

**FIG 2.**
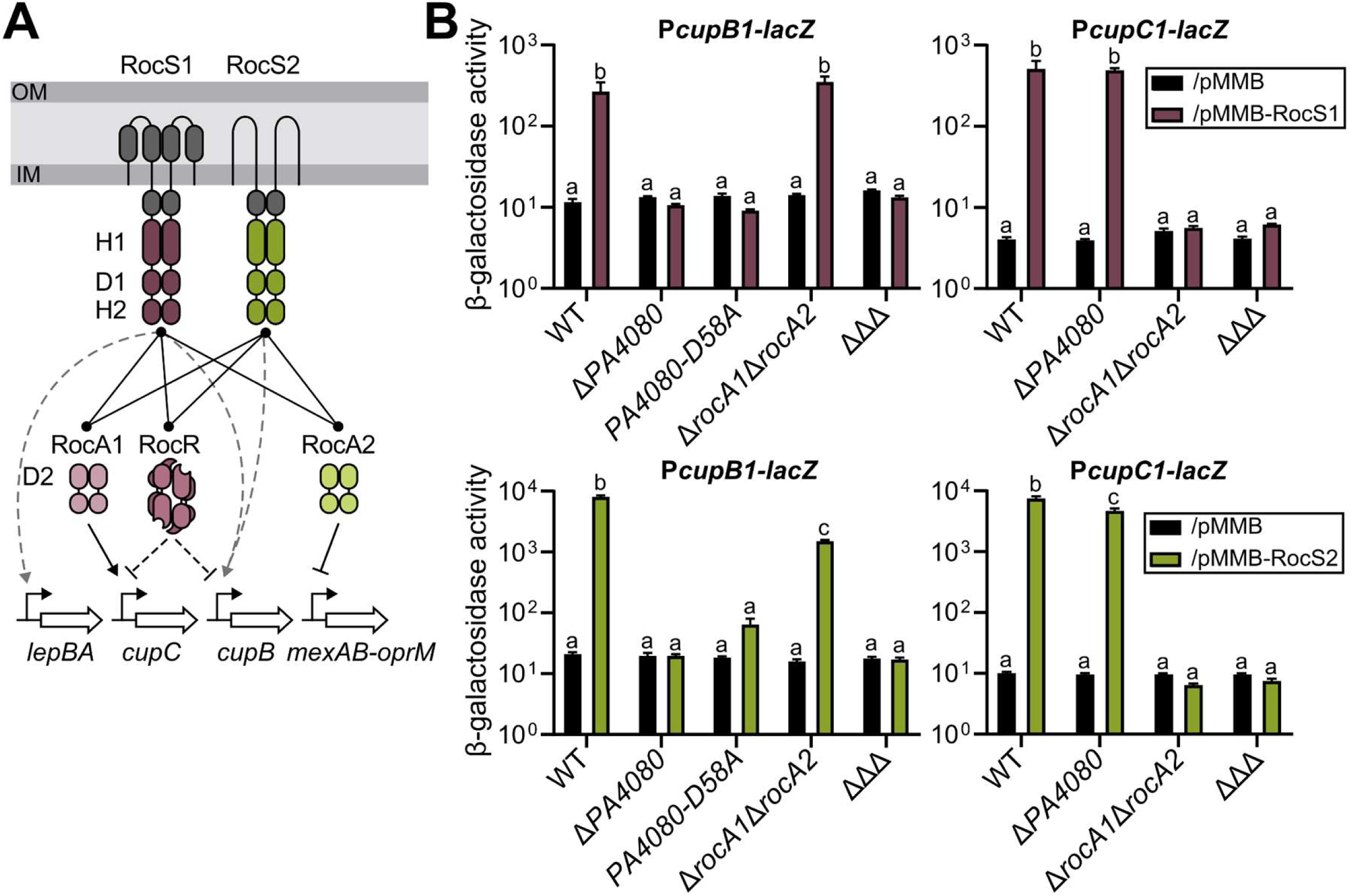
PA4080 is the missing RocA3 regulator. (A) Current view of the Roc signalling pathways. RocR inhibits indirectly expression of *cupC* and *cupB* through c-di-GMPc degradation. Grey dashed lines represent activation through unidentified regulator. RocR is depicted a tetrameric structure as determined by Chen and coworkers (Chen *et al*., 2012). Domains shown are according to PFAM nomenclature; SBP_bac_3 (periplasmic grey domain), PAS_4 (cytosolic grey domain), HisKA and HATPase_c (H1), Response_reg (D1 and D2), Hpt (H2) and either GerE (DNA-binding domain: plain shape) or EAL (c-di-GMP phosphodiesterase; hollow shape) for output domain of RRs. (B) β-galactosidase activities of the strains and mutants carrying the P*cupB1-lacZ* and P*cupC1-lacZ*, as indicated. The strains also carried either the empty pMMB plasmid (black bars), pMMB-RocS1 (burgundy bars) or pMMB-RocS2 (green bars) and the expression of the HKs was induced with IPTG for 6 h in M63 medium. ΔΔΔ corresponds to the triple mutant Δ*rocA1*Δ*rocA2*Δ*PA4080*. The experiments were performed in triplicate and the error bars represent the SEM. Different letters indicate significant differences according to two-way ANOVA followed by Tukey’s multiple comparison test (*p*-value < 0.05).

### PA4080 belongs to the Roc system

To determine whether PA4080 belongs to the Roc system, we assessed its possible activation by the two HKs RocS1 and RocS2 (**Fig. 2A**). To do this, we artificially activated the Roc system by overproducing either RocS1 or RocS2, as previously described (Kulasekara *et al*., 2004; Sivaneson *et al*., 2011). As expected, using chromosomal transcriptional *lacZ* fusions, these overexpressions led to an increased activity of P*cupB1* and P*cupC1* compared to the strains carrying empty vectors in the wild-type genetic background (**Fig. 2B**). The fusions were then introduced into a deletion mutant of the *PA4080* gene, a double mutant Δ*rocA1*Δ*rocA2* and in a strain lacking the three RRs (ΔΔΔ). Deletion of *PA4080* completely abolished the *cupB* operon induction triggered by either HK overproduction, indicating that *cupB* expression is entirely dependent on PA4080 activity and that this RR is the previously unidentified regulator responsible for Roc-dependent regulation of *cupB* (Sivaneson *et al*., 2011). In the Δ*rocA1*Δ*rocA2* mutant, activation of *cupB1* was similar to that observed in the wild-type when *rocS1* was overexpressed, in agreement with previous studies (Kulasekara *et al*., 2004; Sivaneson *et al*., 2011). However, a small but significant reduction in *cupB1* expression was observed upon RocS2 overproduction in the double mutant (**Fig. 2B**), although we know that RocA1 and RocA2 are not direct regulators of the *cupB* operon. These effects, observed only with the HK inducing the highest activities, may result from interruption of autoregulatory loops, as observed for PA4080 (**Fig. 1B**) and RocA1 (Kulasekara *et al*., 2004). Finally, as expected, a complete loss of *cupB* induction was observed in the absence of the three RRs.

We further examined the expression of the *cupC* operon under conditions of RocS1 or RocS2 overproduction in these different mutant backgrounds. Consistent with the absence of control of PA4080 on this promoter (**Fig. S1B**), inactivation of *PA4080* did not affect the activation of *cupC1* expression when *rocS1* was overexpressed. However, we observed a small but significant reduction in *cupC1* expression upon RocS2 overproduction in the *PA4080* mutant, which again may be indirect. To confirm that the *cupB1* induction was due to the activation of PA4080 by HK phosphorylation, a mutation was introduced in *PA4080* to change the conserved aspartate residue of the regulator to an alanine (D58A), preventing RR phosphorylation (**Fig. 2B**). This mutation drastically reduced the activation of *cupB1* expression when RocS1 and RocS2 were overproduced, indicating that this induction requires a phosphorylated PA4080 protein.

Overall, our results indicate that PA4080 is part of the Roc system that mediates the activation of the *cupB* operon by both RocS1 and RocS2 sensory kinases. We have therefore named PA4080 «RocA3».

### RocA1 controls the expression of the *lepBA* operon

In addition to controlling fimbriae and MexAB-OprM synthesis, the Roc system was reported to regulate *lepBA*, as this operon is activated when RocS1 is overproduced. LepBA is a multi-effector secretion system in which LepB transports both the LepA protease and CupB5, encoded in the *cupB* operon, across the outer membrane (Garnett *et al*., 2015). Given this functional link, we suspected that the newly identified RocA3 might be responsible for the activation of *lepBA* expression in addition to that of *cupB*. From the DAP-seq data, we observed that the *lepB* promoter region was indeed targeted *in vitro* by RocA3, but also by the other two RocA proteins, with the higher peak observed for RocA1 (**Fig. 3A**). We therefore investigated the potential regulatory contributions of the three RRs using a P*lepB*-*lacZ* chromosomal transcriptional fusion during activation of the Roc system by *rocS2* overexpression. We first observed a strong activation of *lepB* expression in the wild-type genetic background when RocS2 was overproduced (**Fig. 3B**). Deletion of *rocA1* almost completely abolished this up-regulation suggesting that RocA1 is the main Roc activator of the *lepBA* operon, consistent with the strong DNA binding observed for this RR to the promoter *in vitro* (**Fig. 3A**). Although the effects were small, deletions of *rocA2* and *rocA3* also had a negative effect on the activation of *lepBA* expression by *rocS2* overexpression, which could be explained by their lower binding intensities observed *in vitro* (**Fig. 3A**) or by small indirect effects on RocA1 synthesis or activation.

**FIG 3.**
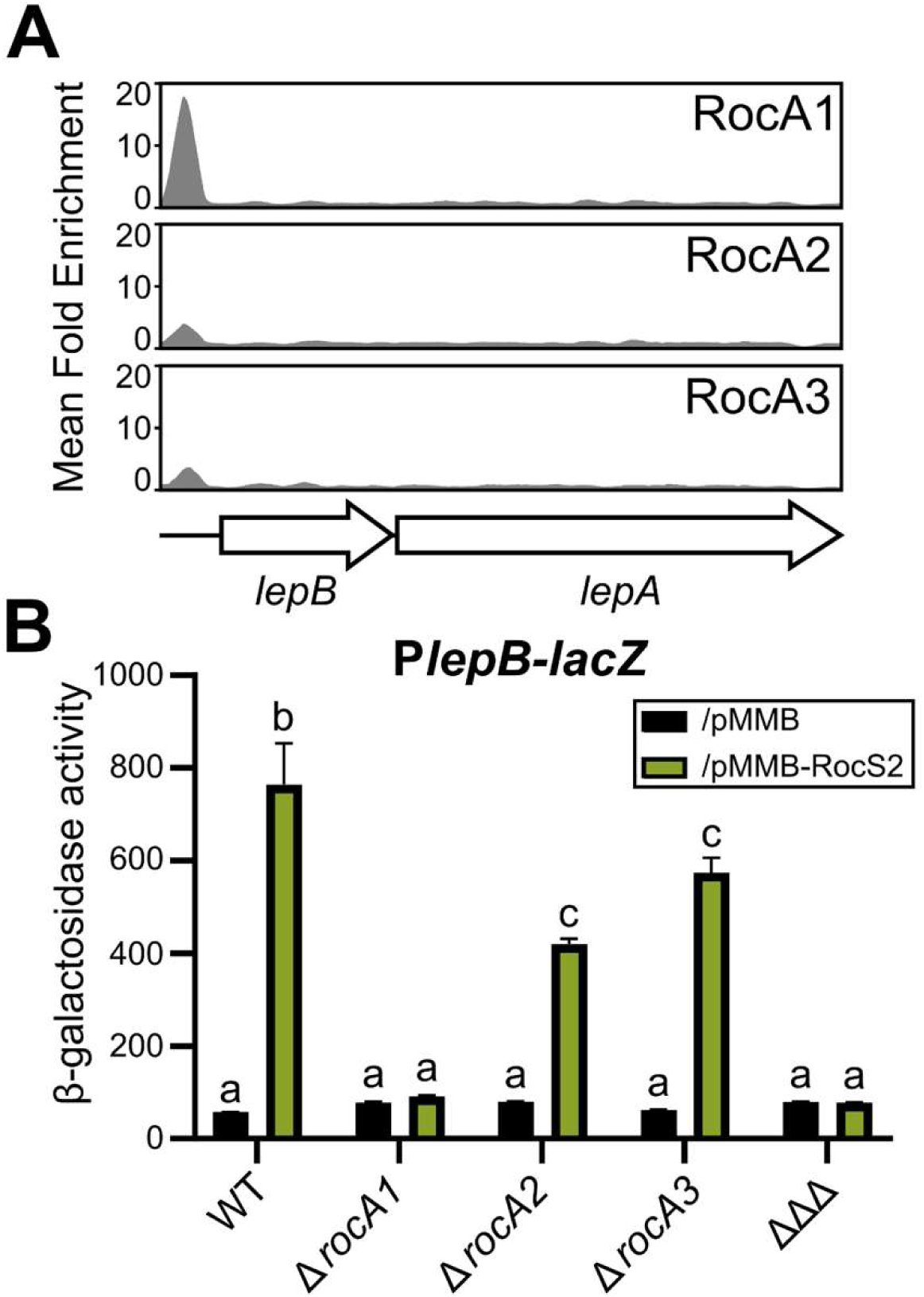
RocA1 controls the synthesis of the LepBA system. (A) Re-analysis of previously published data on the PA14 genome (Trouillon *et al*., 2021) showing the enrichment coverage track of DAP-seq against the negative control for the three RocA regulators on the *lepBA* upstream region. (B) β-galactosidase activities of the strains and mutants carrying the chromosome-integrated P*lepB-lacZ*. Strains also carried either the empty pMMB (black bars) or pMMB-rocS2 (green bars) plasmid and HK expression was induced with IPTG for 6 hours in M63 medium. ΔΔΔ corresponds to the triple mutant Δ*rocA1*Δ*rocA2*Δ*rocA3*. Experiments were performed in triplicate and the error bars represent the SEM. Different letters indicate significant differences according to two-way ANOVA followed by Tukey’s multiple comparison test (*p*-value < 0.05).

Re-analysis of the DAP-seq data (Trouillon *et al*., 2021) enabled us to define the consensus DNA-binding sequences of the three RocA RRs using MEME-ChIP and found them to be highly similar (**Fig. S2A**). Consistent with this observation, we found a large overlap in their putative *in vitro* gene targets, a fraction of which were shared by the three RRs (**Fig. S2BC**); these included the *roc1* locus, the *cupB* and *cupC* operons, and the *lepBA* operon. These targets were shown to be mostly dependent on a single RocA regulator *in vivo* (see above) suggesting that RocA1, RocA2 and RocA3 have specific targets despite their similar DNA-binding sequences. Identified *in vitro* DNA motifs are prone to inaccuracy due to false positives inherent in the method, and discriminating differences between the three RRs may have been overlooked. Of note, RocA2 is the response regulator with the fewest targets, and in the absence of *mexAB-oprM* as a direct target, its mechanism of regulation of the antibiotic resistance phenotype remains unknown.

In conclusion, the Roc system unexpectedly relies on two RRs, RocA1 and RocA3, to orchestrate the expression of the functionally related proteins LepB and CupB5, respectively.

### Organization and diversity of the new *roc3* locus

In PAO1 and PA14, the *rocA3* gene is located adjacent to the *cupB6* gene in a region distinct from the *roc1* and *roc2* loci, which we named the *roc3* locus (**Fig. 4A**). We noted that two additional genes are predicted in this locus, between *cupB6* and *rocA3*, in *P. paraeruginosa* IHMA87 (Winsor *et al*., 2016) (**Fig. 4A**). One is a hypothetical coding sequence (CDS) (*IHMA87*_*RS04355*) that codes for a protein with a predicted Hpt domain, suggesting a potential role in a TCS phosphorelay. The second gene encodes a putative RR (IHMA87_00844, named RocR3 here) that, like RocR, possesses a C-terminal EAL domain that is predicted to have a c-di-GMP-specific phosphodiesterase activity. The *P. aeruginosa* RocR protein has been reported to be a negative regulator of the Roc1 and Roc2 subsystems, inhibiting expression of the *cupB* and *cupC* operons probably indirectly, through degradation of the c-di-GMP second messenger (**Fig. 2A**) (Kulasekara *et al*., 2004; Rao *et al*., 2008; Sivaneson *et al*., 2011). The two proteins share 49% sequence identity and a structural model of RocR3 revealed a similar architecture to that obtained for RocR (Chen *et al*., 2012) (**Fig. 4B**). However, one of the seven conserved residues identified as essential for the catalytic activity of the EAL domain in RocR was not conserved in RocR3 (E357T) (Rao *et al*., 2008) and the motif of the loop 6, which is also involved in the activity (Rao *et al*., 2009), showed two differences (A304S and Y306H), suggesting that the phosphodiesterase activity of the protein might be compromised.

**FIG 4.**
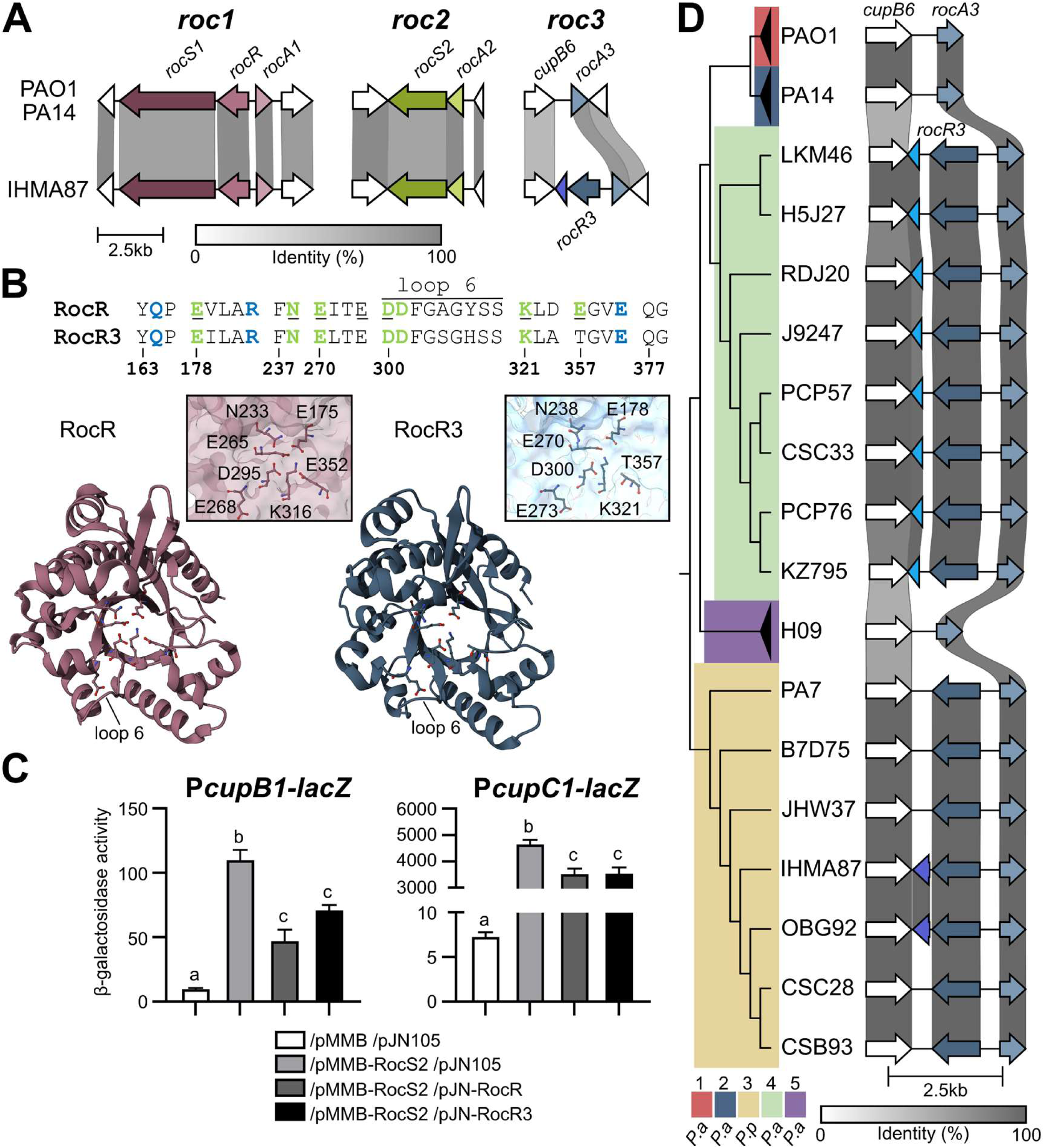
The *roc3* locus in *P. (par)aeruginosa* species. (A) Loci encoding the three Roc systems are represented in the *P. aeruginosa* PAO1 and PA14 strains and *P. paraeruginosa* IHMA87, with the percentage of sequence identity indicated by the grey scale. The genes located on either side of the *roc* genes are also indicated to show the conservation of their location. In the IHMA87 Roc3 system, *rocR3* is homologous to *rocR* while the downstream pseudogene may encode part of an H1 domain, which is an Hpt-like module. (B) Comparison of RocR and RocR3. The upper section shows conserved motifs, with amino acids involved in substrate binding colored in blue and those involved in metal ion coordination colored in green. The seven amino acids identified as essential for catalysis (Rao *et al*., 2009) are underlined (adapted from Romling 2009). The lower section shows an upper view of the structure of the EAL domain of a RocR monomer (3SY8) compared to the model proposed by RoseTTAFold for the EAL domain of RocR3. Details of the organisation of the seven essential residues are shown alongside. The model was obtained using the https://robetta.bakerlab.org/submit.php server, accessed in May 2024 and visualised with Mol* 3D Viewer (Sehnal *et al.,* 2021). (C) β-galactosidase activities of the PAO1 strain containing the P*cupB-lacZ* or P*cupC-lacZ* transcriptional fusion. The strains carried two replicative plasmids as indicated below. Gene expression was induced with 1 mM IPTG and 0.5% arabinose for 6 hours in M63 medium. Experiments were performed in triplicate and the error bars represent the SEM. Different letters indicate significant differences according to two-way ANOVA followed by Tukey’s multiple comparison test (*p*-value < 0.05). (D) Genetic comparison of *roc3* loci in *rocR3*^+^ strains with the percentage of sequence identity indicated by the grey scale. PAO1, PA14 and H09 were used as reference for clades 1, 2 and 5, respectively. Sequences were ordered based on the genomic phylogeny shown.

To assess the role of RocR3 in the Roc network, we compared the effect of RocR and RocR3 overproduction in strains harboring the P*cupB* and P*cupC* transcriptional *lacZ* fusions in the presence of pMMB-RocS2. As expected, RocR overproduction limited significantly the expression of both operons (**Fig. 4C**). Strikingly, the expression of the both operons was also significantly reduced when RocR3 was overproduced, suggesting that the protein is also able to partially antagonize the activation of the RocA1-controlled *cupC* and RocA3-controlled *cupB* operons.

The IHMA87 *roc3* locus is reminiscent of the *roc1* locus in its organization with two divergent genes encoding two RRs, the transcription factor RocA3 and the cyclic diguanylate phosphodiesterase RocR3, except for the absence of *rocS1* (**Fig. 4A**). We examined the conservation of the *roc3* region in 804 complete genomes of *P. aeruginosa* (n=793) and *P. paraeruginosa* (n=7) available in the NCBI genome database. Each genome was grouped into a clade based on phylogeny, using previously established nomenclature (Freschi *et al*., 2019), with the outlier clade 3 now corresponding to *P. paraeruginosa* (Rudra *et al*., 2022) (**Fig. S3**). First, we found that the synteny around *rocR3* was conserved in *P. aeruginosa* strains of clade 4 and in *P. paraeruginosa* strains (**Fig. 4D**). Except for two clade 1 strains carrying a gene encoding a transposase, no other genes were predicted between *cupB6* and *rocA3* in the two main phylogenetic clades (1 and 2) and in clade 5 of *P. aeruginosa*. The adjacent Hpt-encoding gene mentioned above was found in IHMA87 and OBG92 strains but not in other *P. paraeruginosa* strains, although the sequences are highly similar. The prediction of the gene in these two strains results from a 4 bp deletion in the CDS that shifts a stop codon in frame (**Fig. S4**). All clade 4 strains carry a predicted gene next to *rocR3* encoding a HisKA domain that shows strong homology to the H1 domain of RocS1 (**Fig. 4D**). These results led us to the hypothesis that a gene encoding an unorthodox HK (like RocS1) was present downstream of *rocR3* but was lost during evolution. The deletion of this putative *rocS3* gene left different genetic scars between the clades with residual information leading to the prediction of genes coding for parts of this ancestral HK (H1 domain in clade 4 and a Hpt domain reminiscent of an H2 domain in some clade 3 strains) (**Fig. S4**).

In conclusion, an additional RR of the Roc system was identified in the *P. paraeruginosa* IHMA87 strain, RocR3, present in the newly named *roc3* locus which harbors the *rocA3* core gene. The *roc3* locus presents clade-specific gene organization, suggesting gene erosion during species evolution and divergence.

### Synteny of the *cupC* locus suggests additional partners

In the PAO1 and PA14 genomes, upstream of the Roc-regulated *cupC* operon lies a gene encoding one of the three Hpt proteins, HptA (**Fig. 5A**) (Winsor *et al*., 2016). As mentioned above, these Hpt proteins are intermediates for the phosphotransfer between hybrid HKs and their cognate RRs. However, neither one of these proteins is encoded in the vicinity of the *hptA* gene in these two genomes. To assess whether a potential partner of HptA is present in other strains, we used comparative genomics to analyze the genetic environment around this gene in the different *P. aeruginosa* and *P. paraeruginosa* species. In the *P. paraeruginosa* strain IHMA87, we found that a gene (*IHMA87_RS21720*) encoding an orphan hybrid HK is located upstream of the *hptA* gene. The orthologous gene in the genomes of PAO1 (*PA2583*) and PA14 (*PA14*_*30700*) is located elsewhere between two tRNA genes, suggesting that these regions have undergone gene rearrangement (**Fig. 5A**). Like RocS1, PA2583 possesses two predicted periplasmic sensor domains found in solute-binding proteins (SBP_bac_3), which may be involved in ligand binding, and a cytoplasmic PAS domain (Kulasekara *et al*., 2004). Although PA2583 is a hybrid HK and thus differs from RocS1 in the absence of an additional H2 domain, they both share high sequence identities within their H1 and D1 domains (**Fig. 5B**). In addition, their phylogenetic proximity was previously revealed by Chen *et al*. (Chen *et al*., 2004), with PA2583 being the closest HK to RocS1 and RocS2, as mentioned by Sivaneson *et al*. (2011). As the *hptA* gene is contiguous to the gene encoding the predicted hybrid HK PA2583 in IHMA87, we hypothesized that they could be part of a phosphorelay integrated into Roc TCSs. This was supported by the observation that the H2 domain of HptA is very similar to those of RocS1 and RocS2, sharing more than 50% amino acid sequence identity with them. In comparison, the sequence identity of HptB and HptC with the same domains drops below 28% (**Fig. 5B**). Therefore, the sequence similarities and the close localization to the *cupC* operon suggest that PA2583 and HptA could be additional partners of the complex Roc system.

**FIG 5.**
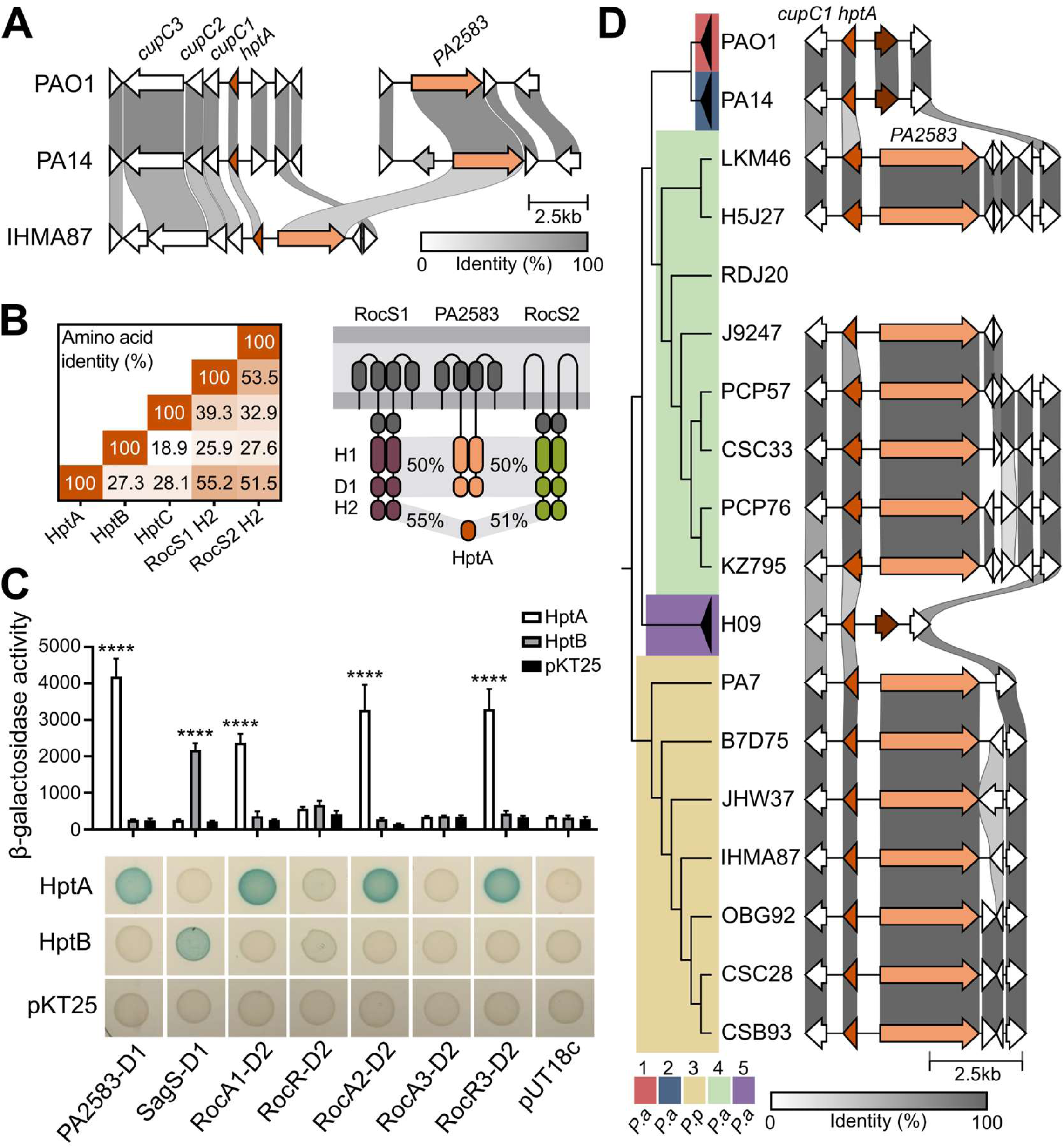
Additional putative players in the Roc system. (A) Genetic organisation of the loci containing the *cupC* operon and the *PA2583* gene in the *P. aeruginosa* PAO1 and PA14 strains and *P. paraeruginosa* IHMA87, with the percentage of sequence identity indicated by the grey scale. (B) The percentage of amino acid identity of the three Hpt modules of PAO1 with the H2 domains of RocS1 and RocS2. The figure on the right shows the homologies of RocS1 and RocS2 unorthodox HKs, PA2583 hybrid HK and HptA proteins. H1, D1 and H2 correspond to the transmitter (HisKA/HATPAse_C), receiver and Hpt domains, respectively. The putative sensory domains are represented by periplasmic grey boxes for the Sbp3 domains and a cytoplasmic grey box for the PAS domain. (C) Bacterial two-hybrid analysis of different combinations of recombinant pKT25 and pUT18c plasmids encoding full or partial proteins of interest in *E. coli* DHM1 spotted on LB agar containing X-Gal and IPTG. Blue colored colonies indicate interacting proteins. β-galactosidase activities of the resuspended spots from bacterial two-hybrid analysis plates are shown on the right. Experiments were performed in triplicate and the error bars represent the SEM. Statistical analysis was performed using two-way ANOVA, followed by Dunnett’s test for comparison with the control condition (pKT25). \*\*\*\**p* < 0.0001. (D) Genetic comparison of the *hptA* locus in strains harbouring PA2583 in the vicinity, with the percentage of sequence identity indicated by the grey scale. PAO1, PA14 and H09 were used as reference for clades 1, 2 and 5, respectively. Sequences were ordered based on the genomic phylogeny shown.

To test this hypothesis, we overproduced the cytosolic part of the PA2583 HK in a strain carrying a transcriptional fusion of the *cupC* promoter with *lacZ*, in combination or not with the overexpression of *hptA*. However, no stimulating effect on *cupC* expression was observed, even when the expression of the entire chromosomal *PA2583* gene was directly driven by a P*BAD* promoter (data not shown). However, we cannot exclude that the overproduced proteins are not active, since no control target was identified. Therefore, we investigated the possible interaction of PA2583 and HptA with different Roc proteins using the bacterial two-hybrid system (Karimova *et al*., 1998) which has previously been used to study interactions in the Roc1 and Roc2 systems (Kulasekara *et al*., 2004; Sivaneson *et al*., 2011). While HptB interacted with its known partner SagS as expected (Hsu *et al*., 2008), we observed that HptA was able to interact with PA2583 and the D2 domains of three RRs, RocA1, RocA2 and RocR3 (**Fig. 5C**), revealing the first identified partners for the HptA protein of *P. aeruginosa*. Therefore, based on sequence homology and interaction analyses, PA2583 was hereafter named “RocS4”.

Considering the different genetic environments of *hptA* in PAO1, PA14 and IHMA87 strains, we further investigated the gene location in *P. aeruginosa* and *P. paraeruginosa* species (**Fig. 5D**). In clades 3 and 4, except for strain RDJ20, *PA2583* was found in the vicinity of *hptA*, whereas it was absent in clades 1, 2 and 5 (**Fig. 5D**). Instead, the gene was always found present in a region surrounded by two tRNAs genes between *PA2584* or *PA2582*. The locus appears to be a hotspot for genetic rearrangements, as the distance between *PA2584* and *PA2582* varies greatly among strains, but *PA2583* was always found in the vicinity of one or the other gene (**Fig S5**). Notably, *PA2583* was predicted to be a pseudogene due to a frameshift in numerous clade 1 and 2 strains.

In conclusion, comparative genomics suggested a possible involvement of a hybrid kinase and the HptA protein in the signaling network of Roc TCSs. Interactions between the HK RocS4 and HptA as well as between this Hpt protein and three RRs were observed *in vivo* suggesting the existence of an additional phosphorelay involving multiple RRs of the Roc system.

### The Roc system contributes to the diversity of the pangenomic repertoire of RRs

Previous studies have established the phylogeny of the RRs present in the *P. aeruginosa* PAO1 strain, but omitted certain RRs such as RocA3 (Chen *et al*., 2004). To integrate RocA3 into the RRs phylogeny and expand our comprehension of the distribution of RRs across the clade, we carried out a genome-wide analysis of RRs at the level of *P. aeruginosa* and *P. paraeruginosa* species. Out of 804 predicted proteomes of the two species, 158 RRs were identified and classified into 21 families according to their domain architectures (**Fig S6A**) (Ortet *et al*., 2015). We then performed a phylogenetic analysis of the REC domain of the 158 RRs protein sequences (**Fig 6**).

**FIG 6.**
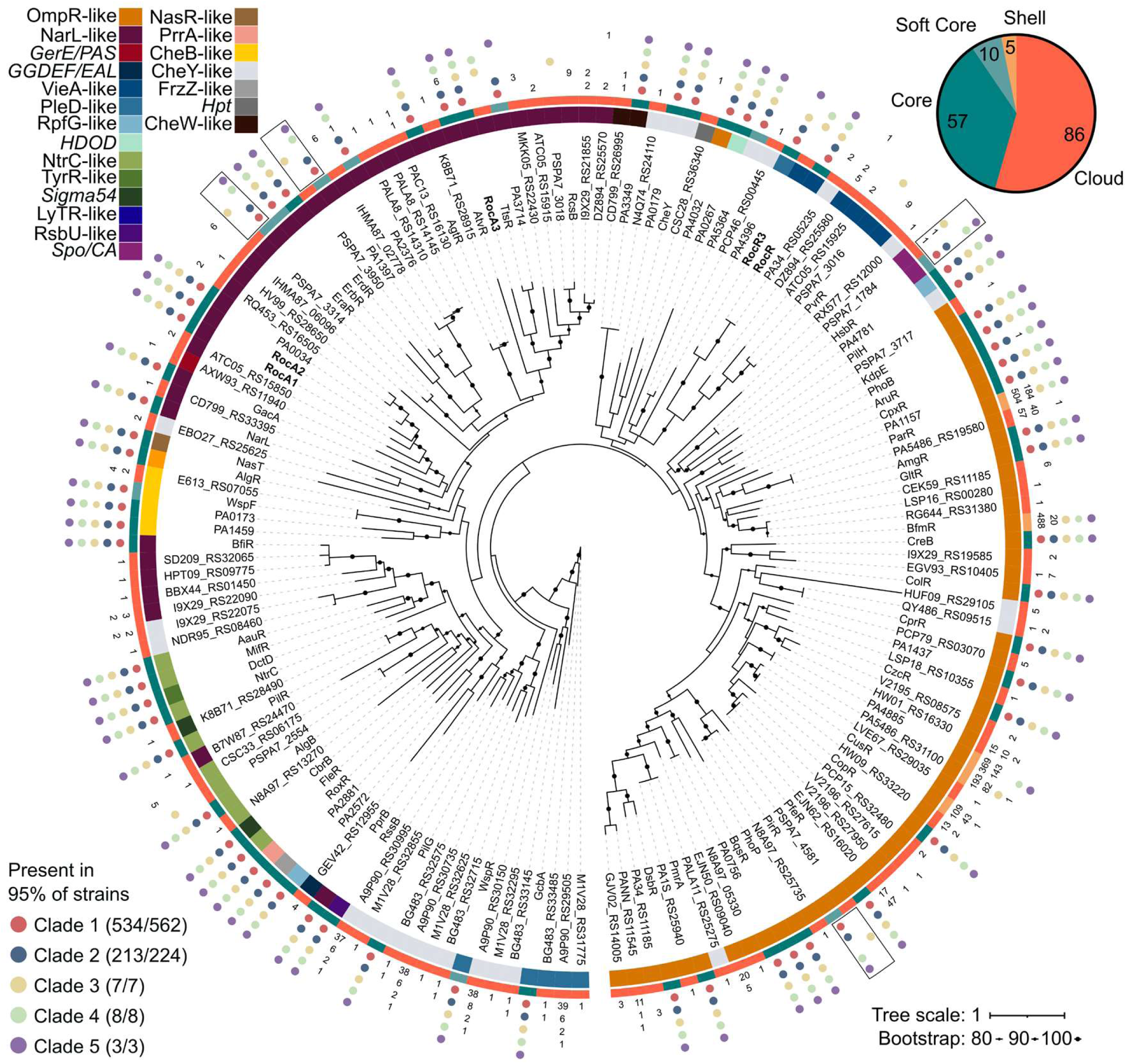
Roc players in the pangenomic repertoire of RRs in *Pseudomonas aeruginosa* and *paraeruginosa*. The unrooted phylogeny of the REC domain of 158 identified RRs is shown in the center. Phylogenetic support is shown by black circles for nodes with high bootstrap value (>80%). Labels correspond to the names of the proteins in PAO1, PA14 or PA7 where possible, otherwise the name of one representant was randomly chosen. Roc actors are highlighted in bold. RRs were classified into families based on their domain architecture, shown by the color in the first ring. The second ring represents whether the gene coding for the protein belongs to the core, soft core, shell or cloud genome of the species. The outer part of the tree shows the conservation of RRs in different clades. Raw counts are given except when more than 95% of the clade harbor the protein, in which case the values are replaced by a colored bubble. The 4 RRs with a specific distribution and separate homologs in clade 3 are shown in boxes.

Our results are broadly consistent with the previous analysis, which grouped the different families together as expected (Chen *et al*., 2004). Proteins with a single REC domain (CheY-like) are found throughout the tree, illustrating the modularity of these proteins, which can lose domains by insertion or recombination. In *P. paraeruginosa*, four RRs have sequences that diverge below the similarity threshold (90% sequence identity) from their homologs in other clades (HsbR, PirR, EraR and PA1397). RRs thus follow different evolutionary rates within the two species studied, suggesting that the redirection of RR functions occurs at different rates. The genome sizes of the strains from the 5 clades are similar with a mean size of 6.74 Mbp, although clade 3 strains which corresponds to the *P. paraeruginosa* species has more RRs with an average of 75 RRs per strains compared to 70 for clades 1, 2, 4 and 5 (**Fig S6BC**). By determining whether these RR-encoding genes belong to the core (genes present in >99% of the strains), soft core (95%), shell (15%) or cloud genome of these species, we evaluated the pangenomic distribution of genes of the different RR families (**Fig S6D**). The most represented families (*ompR*-like, *narL*-like and *cheY*-like) show a broad distribution in the cloud genome, whereas smaller families show a more restricted distribution, some of them exclusively in the core genome (*e. g. nasR*-like). The emergence of new RRs is often the result of duplication, and the restriction of these families to the core genome may be explained by an immediate disadvantage to the appearance of paralogs. The *vieA*-like family is an exception: although poorly represented in the core genome with only *rocR*, genes coding for these RRs are the fourth most abundant in the cloud genome, suggesting that a variable distribution of these regulators between clades is common, as observed for *rocR3*. All strains studied have at least one RR whose gene does not belong to the core or soft core genomes, with clade 3 logically having the most RRs in the shell and cloud genomes (**Fig S5E**).

Concerning the RRs of the Roc system, the genes coding for RocA1, RocA2, RocA3 and RocR are part of the core genome. Within the NarL-like family, RocA3 is phylogenetically distant from RocA1 and RocA2. The latter two are close and are similar to PA0034, which forms a TCS with LadS and regulates *cupA1* (Guo *et al*., 2024). Regarding RocA3, it is closely related to TtsR and PA3714, two homologs separated in our analysis because their level of identity is below the threshold. This separation was observed for the 4 RRs discussed above, but presents a particular pattern for TtsR and PA3714, since TtsR is present in clades 3 and 4. In the strain PA7 of the clade 3, TtsR was shown to regulate expression of *txc*, a secretion system-coding operon located in the RGP69 genomic island inserted just downstream of this RR-coding gene in clades 3, 4 and 5 (Cadoret *et al*., 2014). RocR3 is close to RocR and to a RR with only one REC domain (PA34_RS05235), which results from the insertion of a mobile genetic element into the *rocR* gene in two strains.

In conclusion, the RR regulatory network is largely conserved within *P. aeruginosa* and *P. paraeruginosa* species, which possess 57 core RRs. However, these conserved proteins represent less than half of the diversity of RRs observed at the pangenome scale, demonstrating the high plasticity of these networks. In this study, a clade-specific distribution of RRs was rarely observed and only a few RRs such as PA4396, PSPA7_3016, PSPA7_2554 and RocR3 discriminate the RR composition of the different clades.

## DISCUSSION

Although the Roc system is key to *P. aeruginosa* pathogenicity, our understanding of some of its critical regulatory aspects was still fragmented. This study combined experimental approaches with comparative genomics to unravel the interplay involved in the Roc system, completing the previous model and proposing new partners. Specifically, we characterize four novel players, including two RRs, a hybrid HK and a Hpt protein, almost doubling the size of this interconnected network of TCSs. First, we identified PA4080, which we named RocA3, as a fourth RR capable of being activated by the unorthodox HKs RocS1 and RocS2 and of activating the expression of the *cupB* operon involved in biofilm formation. Although we suspected that RocA3 was the activator of *lepBA*, co-regulating the toxin gene (CupB5) and its transporter (LepB), we found that RocA1 was the main transcriptional regulator of this operon. Comparative genomics allowed us to identify a different genetic organization of the *roc3* locus in *P. paraeruginosa* and in a specific clade of *P. aeruginosa* (clade 4), where a RocR homolog named RocR3 was encoded in the vicinity of *rocA3*. The regulators RocR and RocR3 are VieA-like RRs, both capable of downregulating the expression of the *cupC* and *cupB* operons when overproduced in combination with RocS2. Analysis of the synteny of the *cupC* locus led us to study the hybrid HK PA2583 and the Hpt protein HptA, which showed strong sequence homologies with different domains of RocS1 and RocS2. Genomic comparison revealed that *hptA* is located in close proximity to the RocA1-controlled *cupC* operon in all strains studied. Interestingly, in *P. paraeruginosa* and a specific clade of *P. aeruginosa* (clade 4), *PA2583* is adjacent to *hptA*, whereas the gene is located in a different locus surrounded by two tRNAs in other clades of *P. aeruginosa*. The D1 domain of PA2583 interacts with HptA, which in turns interacts with the D2 domains of RocA1 and RocA2. This is the first report of partners for HptA, which appears to be able to link different RRs of the Roc system to the unorthodox HK PA2583, which we called RocS4 because of its probable implication in the Roc system. The HptA protein was also able to interact with RocR3, highlighting the potential integration of this clade-specific putative RR in the Roc system. Finally, we delineated the repertoire of RR-encoding genes across the pangenome of *P. aeruginosa* and *P. paraeruginosa* and mapped the conservation of these regulators, which can be variable across strains, as shown for the Roc system. Overall, our results have expanded our understanding of the Roc system, proposing a new comprehensive view (**Fig. 7**) by discovering conserved or clade-specific partners that form a plastic and complex network of TCSs that regulate pathogenicity.

**FIG 7.**
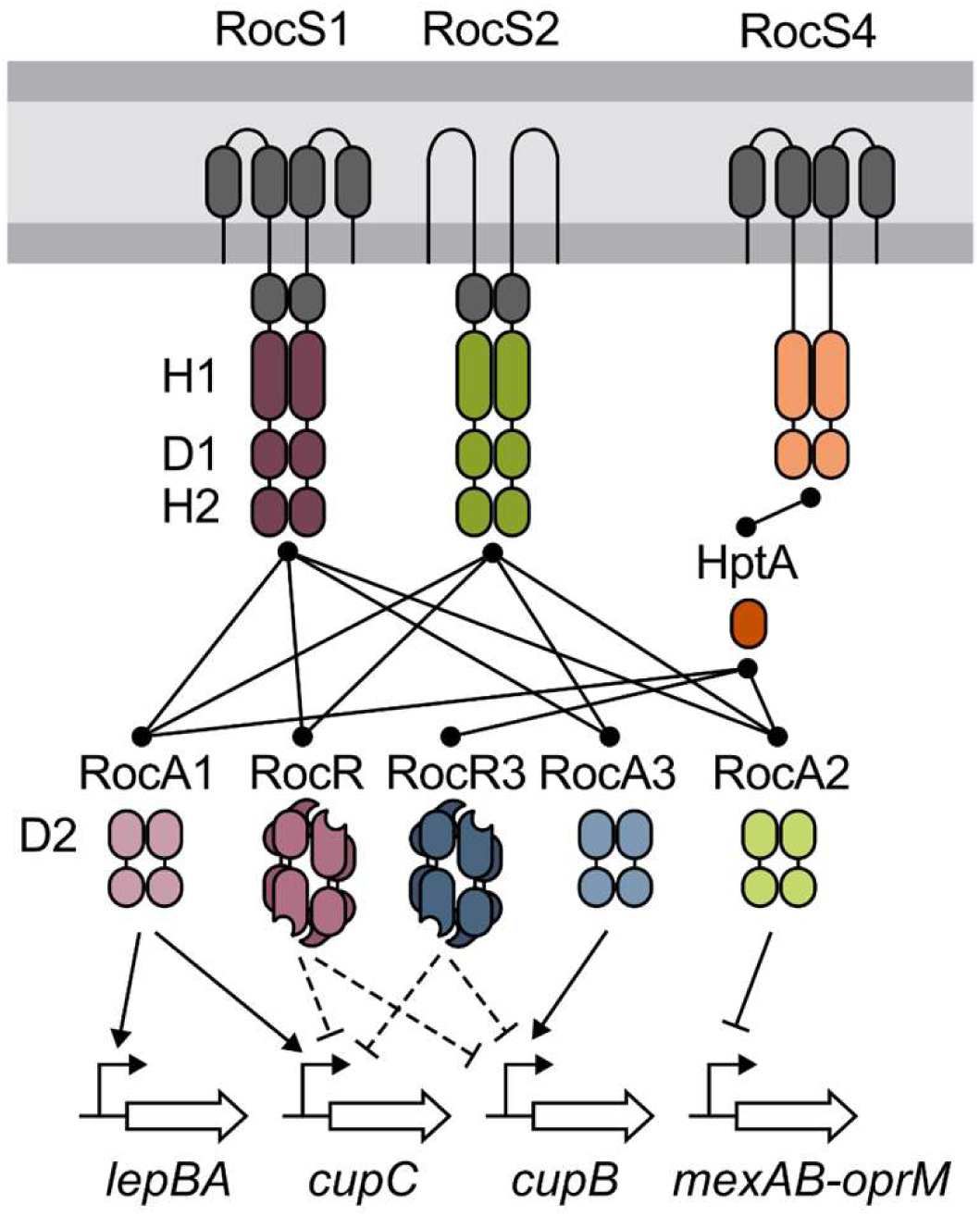
Schematic model of the Roc signaling pathways. RocA3 (PA4080), and the activation of the *lepBA* operon by RocA1 are added. The system was extended with potential new players: the phosphorelay consisting of HptA and the hybrid HK PA2583, named RocS4, and the c-di-GMP specific phosphodiesterase RocR3 found in a few *P. aeruginosa* and *P. paraeruginosa* strains. RocR3 is depicted with the same tetrameric structure as RocR (Chen *et al*. 2012). Domains shown are according to PFAM nomenclature; SBP_bac_3 (periplasmic grey domain), PAS_4 (cytosolic grey domain), HisKA and HATPase_c (H1), Response_reg (D1 and D2), Hpt (H2) and either GerE (DNA-binding domain: plain shape) or EAL (c-di-GMP phosphodiesterase; hollow shape) for output domain of RRs.

In Roc signaling pathways, the HKs RocS1 and RocS2 are able to interact and activate multiple RRs, whose genes are located in different loci. These HKs possess different sensory domains, potentially offering the integration of different incoming signals. Indeed, as predicted for the hybrid HK PA2583, RocS1 possesses two consecutive solute binding protein (SBP) domains belonging to the SBP_bac_3 (PF00497) family, which are known to frequently detect amino acids (Matilla *et al*., 2021). In addition, the three HKs possess a cytoplasmic PAS (Per-Arnt-Sim) domain that could be involved in protein dimerization, ligand binding or oxygen/redox sensing (Henry and Crosson, 2011). Because the Roc system is able to reduce the antibiotic resistance of the bacteria while activating their biofilm formation, it was suggested that it may be important for the adaptation to the lungs of cystic fibrosis (CF) patients (Sivaneson *et al*., 2011). The signals perceived by HKs are rarely known and the study of TCSs often involves their artificial activation, which can be achieved by overexpressing the RR or HK genes, as was done in this work. However, this is not always effective, as observed with PA2583, and predicting the outcome of such overexpression is difficult. Indeed, competition between different TCS partners involves finely tuned concentrations of HKs and RRs for a proper signaling. For example, HKs are usually less numerous than RRs to favor out-competition within RRs and reduce cross-talks with noncognate RRs (Laub and Goulian, 2007; Siryaporn and Goulian, 2008). In addition, phosphotransfer efficiency can differ between an HK and its multiple partners as shown for the asymmetric cross-talk between NarXL and NarQP, with NarX interacting preferentially with NarL (Noriega *et al*., 2010). Finally, HKs can be bifunctional, controlling both phosphorylation and dephosphorylation of cognate RR(s). Therefore, artificial activation of a system may lead to hazardous results that must be distinguished from natural *in vivo* activation. In the Roc system, we do not know if RocS1 and RocS2 are bifunctional HKs. Moreover, their overproduction activates the signaling pathways only in M63 medium and not in LB, suggesting a potential lock that is lifted in the favorable medium. Only the activation of the HKs by their physiological signals could allow the normal functioning of the systems, integrating all the dynamics of the different partners and allowing us to determine the exact extent of cross-talk and cross-regulation.

Why should bacteria maintain all the interconnected TCSs of the Roc system? When TCSs are duplicated, the proteins usually need to rapidly gain new functions in order to be maintained through different mechanisms, such as changes in HK sensory domains and pathway inputs or changes in RRs and pathway outputs (Capra and Laub, 2012). As mentioned above, RocS1 has specific periplasmic sensory domains that its closest paralog RocS2 does not possess, suggesting that they have undergone rearrangement of their sensory domains. Furthermore, although the identified binding sites of RocA1, RocA2 and RocA3 present similarities, they control different targets *in vivo*, indicating that their transcriptional activity differs (binding capability and/or interaction with RNA polymerase). In conclusion, although communication within the network formed by the Roc partners appears to be interconnected, it seems that the signal detection domains of the HKs and the effector domains of the RRs are very specific to each component, allowing multiple genes to be controlled in response to different potential signals. The rationale behind the interconnection of TCSs remains elusive, the simplest explanation being the ability to integrate multiple stimuli to regulate a large number of genes. Such a network needs to be considered in the light of physiological activation implementing, for example, competition between RRs and feedback regulatory loops, as seen here with RocA3, making the Roc system probably a more subtle system than just a relay between multiple signal and multiple regulators.

Our RRs phylogeny raises questions about the Roc system and its variability, which seem to escape every simple evolutionary scenario. The phylogenetic distance observed between RocA3 and RocA1/RocA2 is unexpected in two respects. Functionally, our study shows that RocA3 is integrated into the Roc system, demonstrating that, despite differences, this RR is able to communicate with RocS1 and RocS2. In addition, RocR3, whose gene is located in the same locus as *rocA3* in some strains, is phylogenetically very close to RocR, suggesting a duplication. Furthermore, the polymorphisms observed at the *roc3* and *hptA* loci are surprisingly incongruent with the established species tree in the way that clades 1, 2 and 5 are grouped together. The explanation for this discrepancy is not clear, since the genetic scars left by the putative loss of *rocR3* and *PA2583*, respectively, are similar in these clades, tending to exclude the parallel loss and rearrangement of the genes (**Fig. S4**). It is then tempting to assume that the maintenance of the ancestral polymorphism observed in the intermediate clade 4 for both loci arose from deep coalescence, providing a new perspective on our understanding of the evolution of *P. aeruginosa* and *P. paraeruginosa* species. Finally, the reason for these different organizations remains elusive, and one can only speculate on the functioning of the system in these clades based on our observation for the PAO1 strain.

In conclusion, TCS conservation within species is variable and, as we showed in *P. aeruginosa* and *P. paraeruginosa*, the distribution and organisation of the genes encoding these regulatory proteins are important determinants of the strain-dependent diversity of TCS regulatory networks (Trouillon *et al*., 2021; Elsen *et al*., 2024). Our work showed how a one-of-a kind TCS network, mostly conserved in *P. aeruginosa* and *P. paraeruginosa* species, was differentially shaped in these species by loss and recruitment of different players.

## MATERIALS AND METHODS

### Bacterial strains and growth conditions

The *Escherichia coli* and *P. aeruginosa* strains used in this study are described in Table S1. Bacteria were grown aerobically in lysogeny broth (LB) or in M63 minimal medium (100 mM KH_2_PO_4_, 15 mM (NH_4_)_2_SO_4_, 1.7 µM FeSO_4_, 1 mM MgSO_4_, 0.2% glucose, 0.5% casamino acids, pH7) at 37°C. *P. aeruginosa* was also cultured on Pseudomonas Isolation Agar plates (PIA Difco). To assess the β-galactosidase activities of strains carrying chromosome-integrated *lacZ* fusions, LB-grown cells were diluted to OD_600_ of 0.1 in M63 medium with the appropriate antibiotics and inducers (0.5% arabinose for pJN105-derived plasmids; 10 µM IPTG for pMMB-rocS1 and 1 mM IPTG for pMMB-rocS2) when required and incubated with shaking for 6 hours before assays were carried out. Antibiotics were added at the following concentrations (in µg/ml): 100 ampicillin (Ap), 10 chloramphenicol (Cm), 50 gentamicin (Gm), 10 tetracycline (Tc) for *E. coli*; 300 carbenicillin (Cb), 200 Gm, 200 Tc on PIA plates, 200 Cb and 120 Gm in LB liquid culture for *P. aeruginosa*.

### Plasmids and genetic manipulation

The plasmids utilized in this study and the primers used for PCR are listed in Tables S1 and S2, respectively. All the constructions were verified by sequencing.

To generate *P. aeruginosa* deletion mutants, upstream (sF1/sR1) and downstream (sF2/sR2) flanking regions of the different genes were fused and cloned into the *Sma*I-cut pEXG2 plasmid by sequence- and ligation-independent cloning (SLIC) using the appropriate primer pairs (Li and Elledge, 2007). The same strategy was used to create the point mutation within the *rocA3* sequence, with the overlapping primers creating the mutation. To create the transcriptional *lacZ* fusions, fragments comprising the ATG and around 500 pb upstream of each analysed target were amplified using the appropriate pairs (sF/sR) and inserted into *Sma*I-cut miniCTX-TrrnB-lacZ by SLIC. To express genes under the P*BAD* promoter, sequences containing the entire coding sequences with 50 bp upstream were amplified using the appropriate pairs (sF/sR) and inserted into *Sma*I-cut pJN105 by SLIC. For the plasmids used in the bacterial two-hybrid systems, the sequences were amplified using the appropriate pairs (sF/sR) and inserted into *Bam*HI-cut pKT25 or pUT18c by SLIC: this created hybrid proteins with heterologous proteins fused to the C-terminal of T25 or T18 fragment of the *Bordetella pertussis* adenylate cyclase, respectively.

The pJN105- and pMMB-derived plasmids were introduced in *P. aeruginosa* by transformation while the pEXG2- and miniCTX-TrrnB-*lacZ*-derived vectors were transferred into *P. aeruginosa* strains by triparental mating using the helper plasmid pRK600. Allelic exchanges for mutagenesis were selected as previously described (Berry *et al*., 2018). To excise the miniCTX backbone of strains carrying the *lacZ* fusions, the pFLP2 plasmid was first introduced in the cells and, in a second step, the bacteria were streaked on medium containing 5% sucrose to select for the loss of pFLP2. To create the mutant encoding the RocA3_D58A_ protein, the mutated sequence was introduced in the Δ*rocA3* mutants to replace the deleted gene.

### β-galactosidase activity assay

β-galactosidase activity was assayed as previously described (Miller, 1977; Thibault *et al*., 2009), at least in triplicate. Activities were expressed in Miller Unit (MU) and the error bars in the graphs indicate the standard error of the mean (SEM). When relevant, statistical significance was assessed using the ANOVA test with Tukey’s or Dunnett’s method.

### Bacterial two-hybrid assay

This assay was conducted as previously described (Kulasekara *et al*., 2004). The nomenclature used to define the proteins was as follows: the transmitter domain of the HK carrying the conserved histidine residue is called H1. Unorthodox HKs possess additional domains called D1 and H2. The receiver domain of the RR carrying the conserved aspartate residue is the domain D2. The plasmids pKT25-HptB and pUT18c-SagS-D1 were used as positive controls. The pKT25- and pUT18c-derived plasmids were co-transformed into the DHM1 strain using heat shock and single colonies were patched on LB containing 5-bromo-4-chloro-3-indolyl-beta-D-galactopyranoside (X-gal) at 40 µg/ml and 1 mM IPTG. Positive interactions were identified as blue colonies on X-gal after 24 h incubation at 30°C. Colonies were resuspended in water for β-galactosidase assays.

### DAP-seq analysis

Published DAP-seq data were collected with GEO Series accession number GSE179001 (Trouillon *et al*., 2021). DNA-binding motif discovery was performed using MEME-ChIP (Machanick and Bailey, 2011). Relative fold-enrichments were calculated for each RR on each genome. The PA14 dataset was used based on quality thresholds because the PA4080 experiment with the PAO1 genome didn’t provide sufficient DNA enrichment.

### Phylogenetic analyses

Phylogenetic analyses of *P. aeruginosa* and *P. paraeruginosa* were performed on 804 annotated genomes available in NCBI and Pseudomonas.com databases assessed on April 2024. Core gene alignments were obtained from Roary (minimum 90% identity and a presence in 99% of isolates to be considered core) (Page *et al*., 2015), and SNPs were extracted using SNP-sites (Page *et al*., 2016). For phylogenetic analyses of RRs, 158 RRs were identified by determining the domain architecture of the proteins in the pangenome obtained from Roary with InterProScan (Jones *et al*., 2014), and REC domains were aligned with MAFFT with the L-INS-I option (Katoh and Standley, 2013). Maximum-likelihood trees were build using FASTTREE for genomes (Price *et al*., 2010) with default parameters and IQ-tree with 1000 bootstrap replicates for REC domains (Nguyen *et al*., 2015), then visualized and annotated using iTOL (Letunic and Bork, 2021).

### Bioinformatic analyses

Pangenomic distributions of genes were obtained from Roary. Sequence alignment and visualization were performed using Clinker (Gilchrist and Chooi, 2021). Sequence homologies were evaluated with Clustal Omega (Madeira *et al*., 2022). Variation in genetic regions was examined using Jalview (Waterhouse *et al*., 2009). Most tools were used on the European or American Galaxy servers (The Galaxy Community *et al*., 2022).

## ACKNOWLEDGMENTS

We thank Alain Filloux for providing the pMMB67EH, pMMB67-RocS1 and pMMB67-RocS2 plasmids and Christophe Bordi for the pUT18c-RocA1-D2, pUT18c-RocA2-D2 and pUT18c-RocR-D2 plasmids. We would also like to thank Sophie Abby for her advice and helpful discussions. This work was supported by GRAL, funded through the University Grenoble, Alpes Graduate School (Ecoles Universitaires de Recherche) CBH-EUR-GS (ANR-17-EURE-0003). We also acknowledge the support of the CNRS, the CEA and the Grenoble Alpes University. The Ph.D. fellowship for VS was funded by the French Ministry of Education and Research.

## AUTHOR CONTRIBUTIONS

**Victor Simon:** Conceptualization; investigation; formal analysis; writing – original draft; writing – review and editing. **Julian Trouillon:** Formal analysis; writing – review and editing. **Ina Attrée:**

Writing – review and editing. **Sylvie Elsen:** Conceptualization; investigation; writing – original draft; writing – review and editing.

## DATA AVAILABILITY STATEMENT

The data that support the findings of this study are available in the supplementary material of this article.

## ABBREVIATED SUMMARY

The Roc system represents a highly interconnected but incomplete network of two-component regulatory systems involved in the virulence of *Pseudomonas aeruginosa*. Our work has enabled us to identify the missing RocA3 regulator and to propose new players in the system and to delineate their conservation between clades of the species.

---

**FIG S1.**
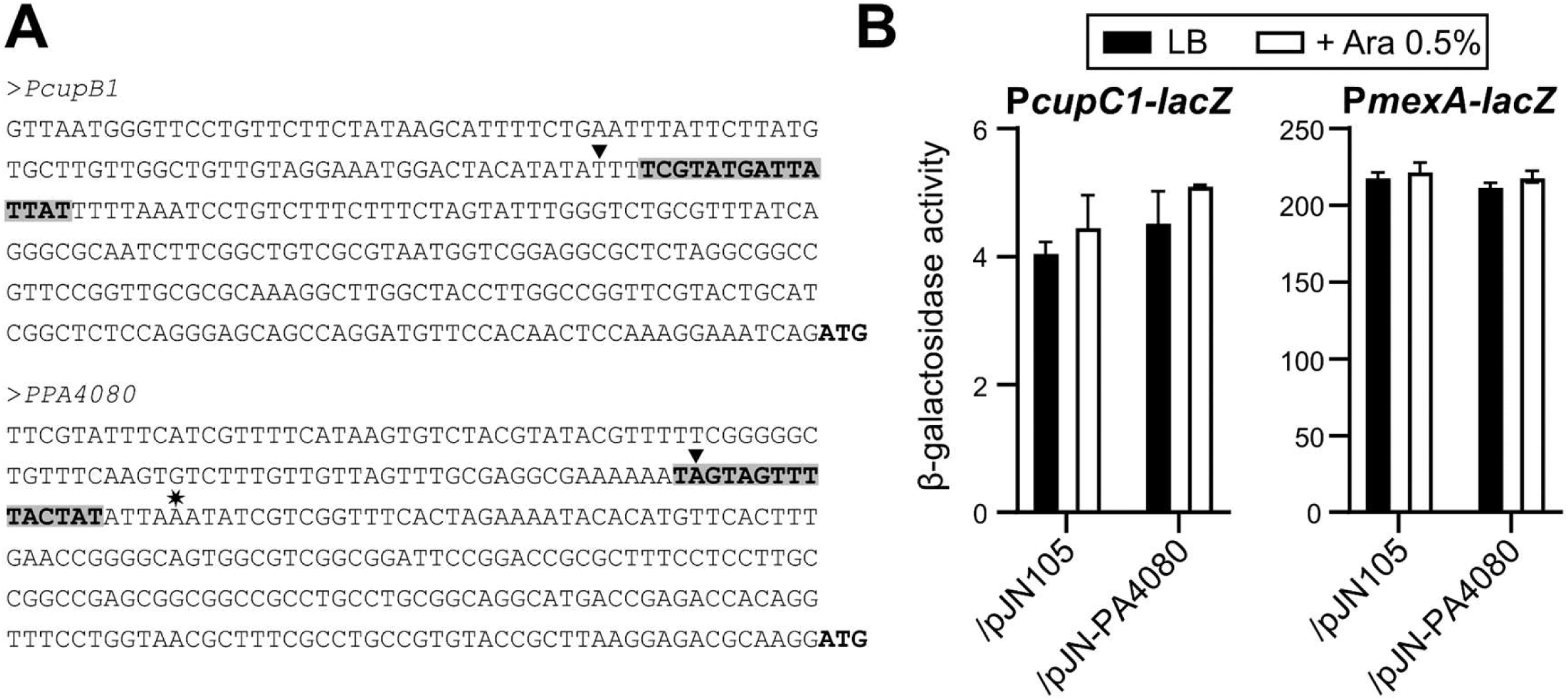
PA4080 regulates the *cupB* operon, its own expression but not other Roc targets. (A) Upstream sequences of the *cupB1* and *PA4080* genes. The ATG of each coding sequence is indicated as well as the location of the transcriptional start site of *PA4080* (star). The predicted binding site for PA4080 on each sequence is shown in bold and colored in grey, with the top of the DAP-seq peak (Trouillon *et al*., 2021) indicated by an arrowhead. (B) β-galactosidase activities of the indicated strains carrying the P*cupC1-lacZ* or P*mexA-lacZ* transcriptional fusions. The strains also carried either the empty pJN105 or the pJN-PA4080 plasmid, and expression of *PA4080* was induced with 0.5% arabinose for 2.5 h in LB medium. Experiments were performed in triplicate and the error bars represent the SEM. Statistical analysis was performed using two-way ANOVA, followed by Dunnett’s test for comparison to the control condition (PAO1 WT /pJN105 in LB). \*\*\*\**p* < 0.0001.

**Fig S2.**
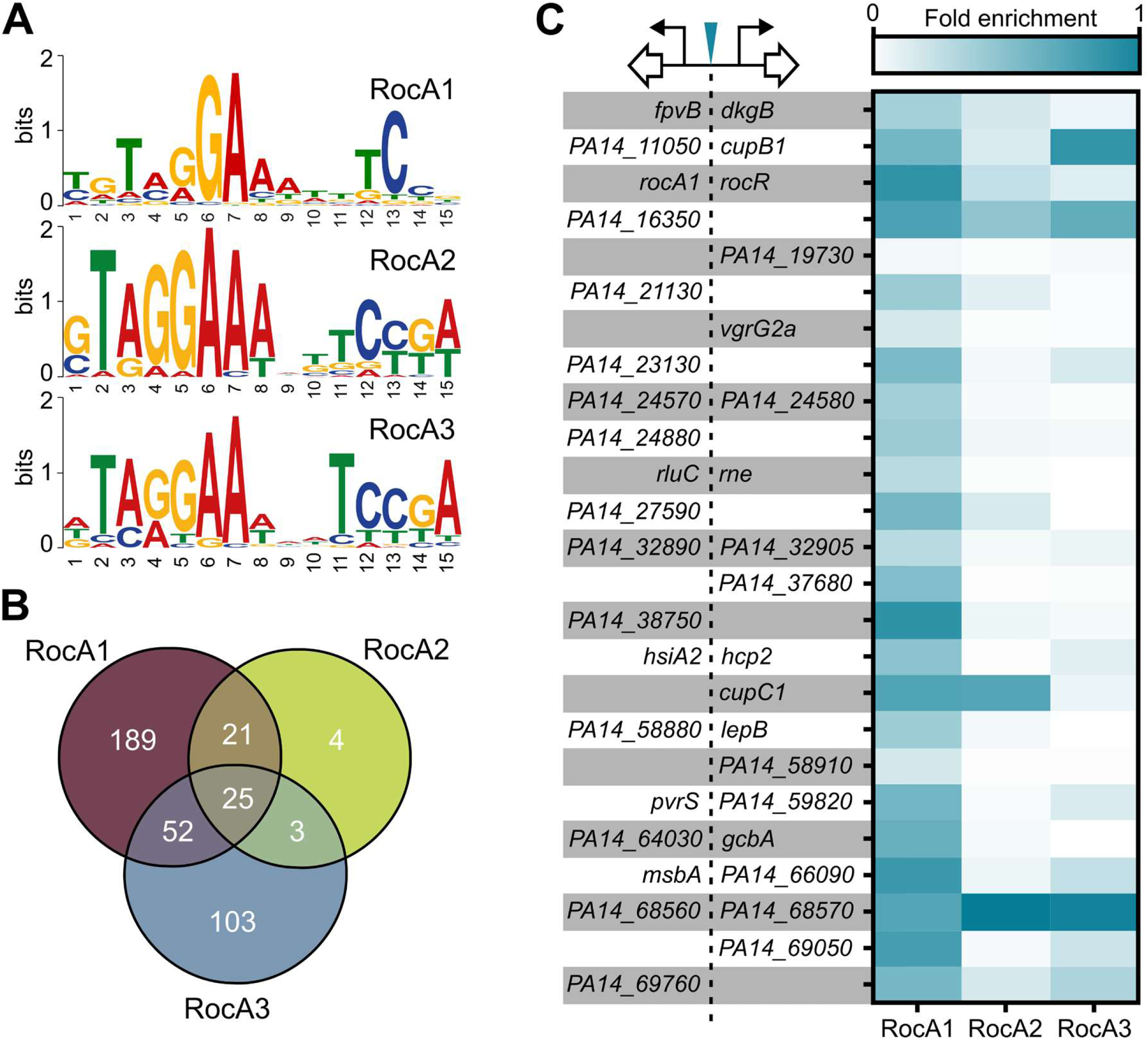
RocA1, RocA2 and RocA3 targets on the PA14 genome. Reanalysis of the dataset of DAP-seq results from published data (Trouillon *et al*., 2021) (A) DNA-binding motifs of RocA1, RocA2 and RocA3 identified by MEME-ChiP. The amino acid symbols are shown at each position. Sequence conservation at each position is indicated by the total height of the stack. The relative frequency of each amino acid is indicated by the height of the symbols within the stack. (B) Overlap of the inferred targets of RocA1, RocA2 and RocA3. (C) Relative fold-enrichment of 25 common targets of RocA1, RocA2 and RocA3. Intergenic target regions are represented on the left, indicating the first gene of potentially regulated transcription units.

**FIG S3.**
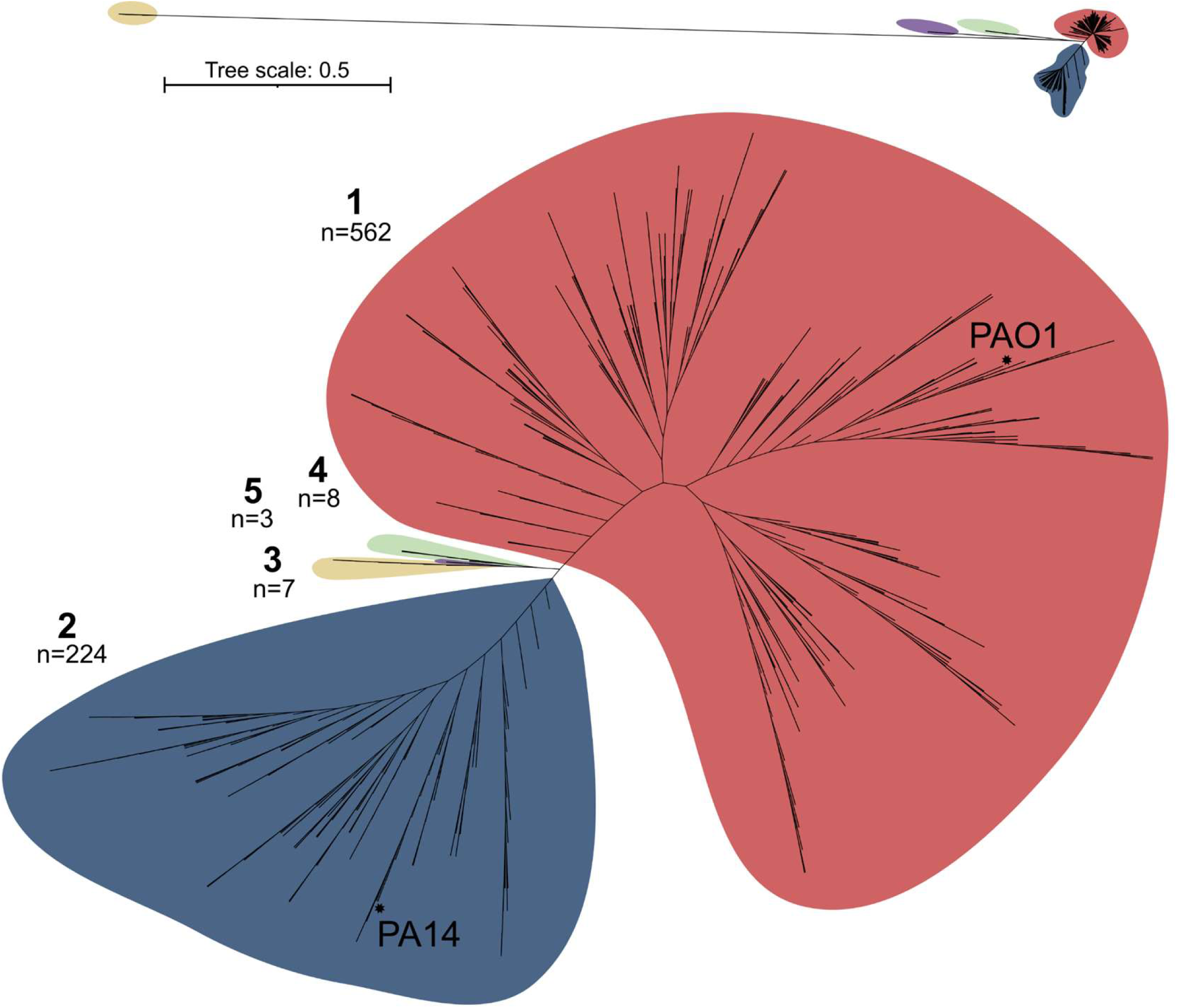
Unrooted phylogenetic tree of *P. aeruginosa* and *P. paraeruginosa* species. The number of genomes for each clade is given (n=804) and the positions of the PAO1 and PA14 reference strains are indicated by a star. The small tree in the upper part represents actual genetic distances between the clades.

**Fig S4.**
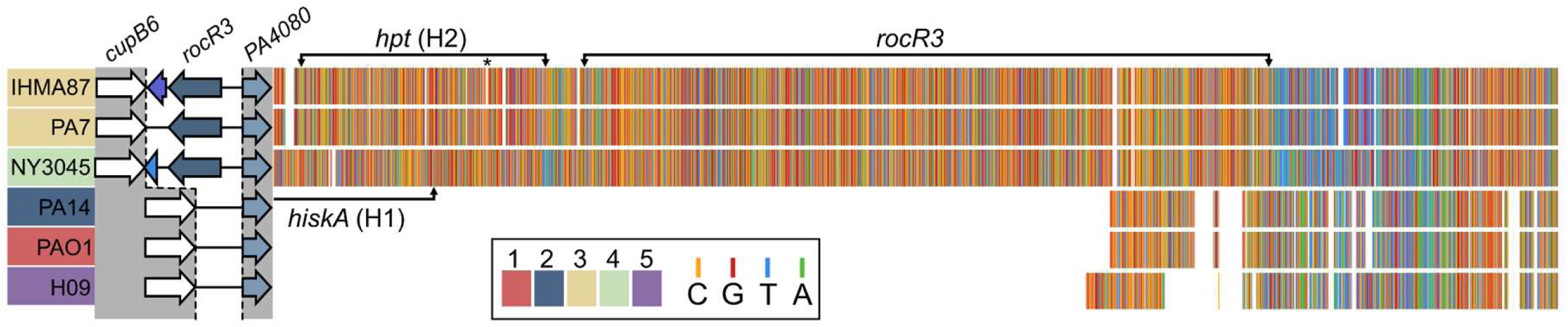
Genetic variations in the *roc3* locus. Alignment of 6 sequences of the genetic region between *rocA3* and *cupB6* for six strains from different clades. The position of different CDSs is highlighted by brackets: the Hpt-encoding gene (*hpt* or H1 domain), the HiskA-encoding gene (*hisKA* or H2 domain), and *rocR3*. The asterisk indicates the 4-bp deletion in IHMA87.

**Fig S5.**
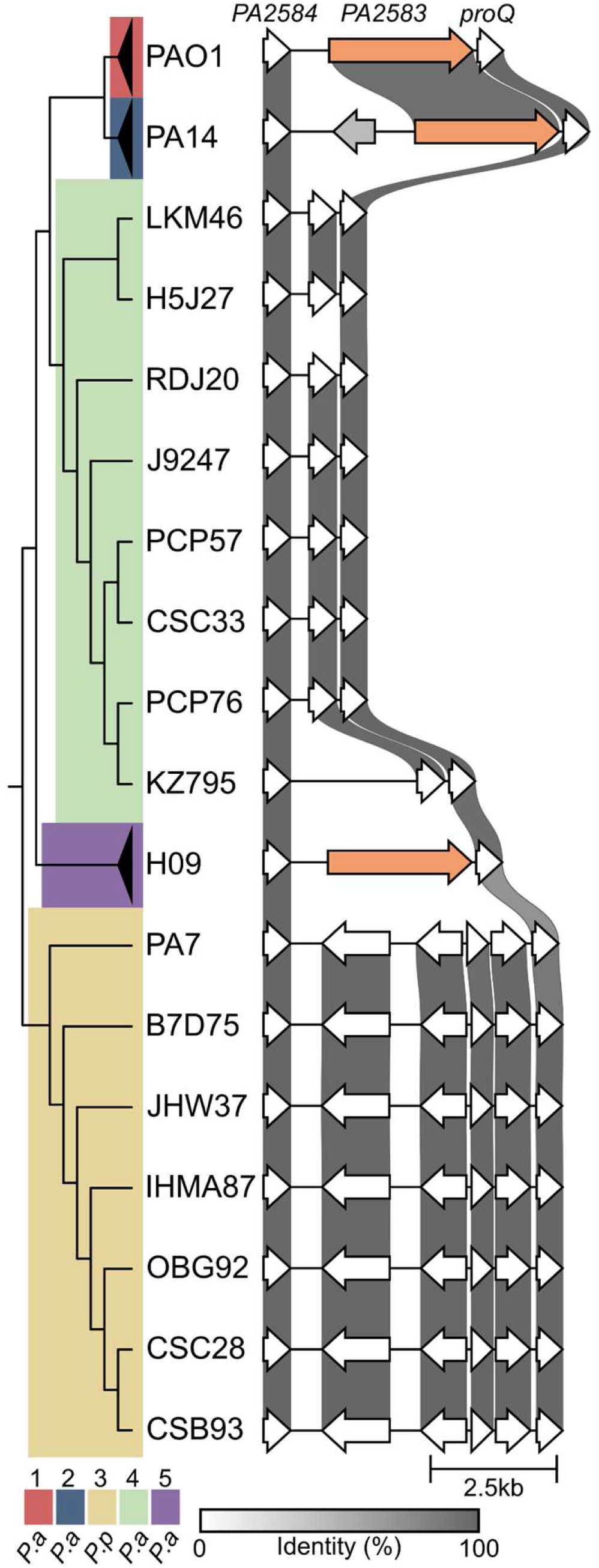
Genetic comparison of the *PA2583* locus in clade 3 and 4 strains, with the percentage of sequence identity indicated by the grey scale. PAO1, PA14 and H09 were used as reference for clades 1, 2 and 5, respectively. Sequences were ordered based on the genomic phylogeny shown.

**Fig S6.**
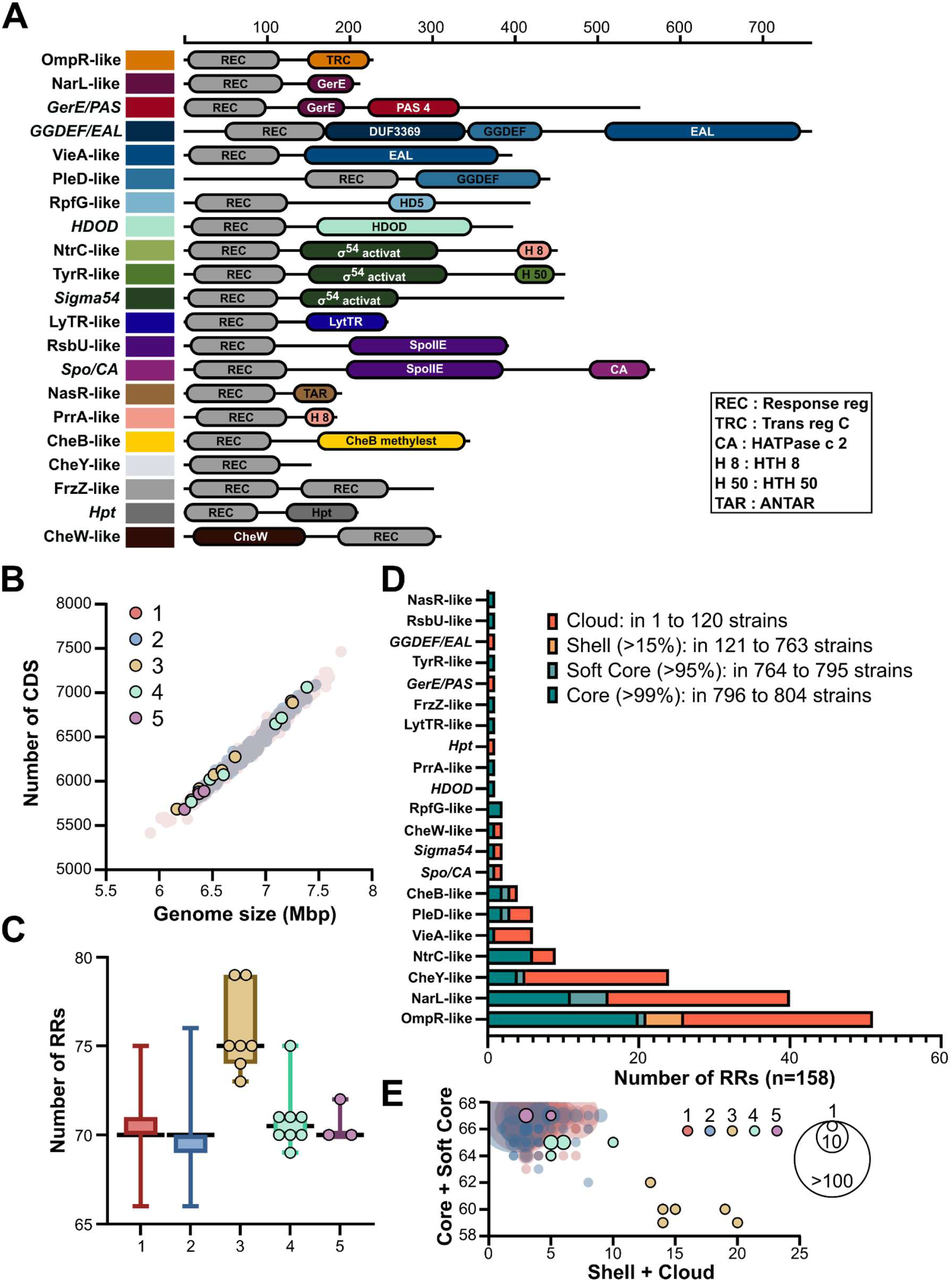
Pangenomic repertoire of RRs in *Pseudomonas aeruginosa* and *Pseudomonas paraeruginosa*. (A) Domain architectures of the different RRs families identified using Pfam names. The length of the domains corresponds to the average of the different representatives for each family. The names of the families are chosen according to previous work (Ortet et al., 2015), except for six families whose names, in italics, were given based on the domain associated with the REC domain. (B) Genome sizes and total number of predicted CDSs for the 804 genomes of *P. aeruginosa* and *P*. *paraeruginosa* classified by clades. Superabundant representatives of clades 1 and 2 are shown as transparent. (C) Number of RRs identified in each genome classified by clade. The boxes represent the mean, maximum, and minimum values, as well as the 95% interval of the distribution. Individual values are shown for clades 3, 4 and 5. (D) Distribution of different RRs in the core, soft core, shell and cloud genomes classified into families. (E) Distribution between core/soft core RRs and shell/cloud RRs according to different genomes. The size of the bubble reflects the number of genomes with identical values. Clades 1 and 2 are shown as transparent.

