## Supplemental Tables S1 & S2 for "Functional and pangenomic exploration of Roc two-component regulatory systems identifies novel players across *Pseudomonas* species"

**Table S1. Bacterial strains and plasmids used in this work**

| **Strain or plasmid** | **Genotype or relevant properties** | **Reference/Source** |
| --- | --- | --- |
| **Strains** | | |
| *P. aeruginosa* | | |
| PAO1 | Wound isolate, sequenced laboratory strain | J. Mougous |
| PAO1Δ*PA4080* | PAO1 with *PA4080* (*rocA3*) deletion | This study |
| PAO1Δ*rocA1* | PAO1 with *PA3948* (*rocA1*) deletion | This study |
| PAO1Δ*rocA2* | PAO1 with *PA3045* (*rocA2*) deletion | This study |
| PAO1Δ*rocA1*Δ*rocA2* | PAO1 with *rocA1* and *rocA2* deletion | This study |
| PAO1Δ*rocA1*Δ*rocA2*Δ*rocA3* | PAO1 with *rocA1*, *rocA2* and *rocA3* deletion | This study |
| PAO1-PA4080 D58A | PAO1 with *rocA3* mutated gene encoding RocA3D58A | This study |
| ***P. paraeruginosa*** | | |
| IHMA87 | wild-type strain (urinary infection) | (Kos *et al.*, 2015) |
| *E. coli* | | |
| TOP10 | Chemically competent cell | Invitrogen |
| DHM1 | F- *cya-854 recA1 endA1 gyrA96* (NaIR) *thi1 hsdR17 spoT1 rfbD1 glnV44*(AS) | (Karimova *et al.*, 1998) |
| **Plasmids** | | |
| pRK600 | Helper plasmid with conjugative properties (CmR) | (Kessler *et al.*, 1992) |
| pFLP2 | Plasmid expressing the Flp recombinase (ApR/CbR) | (Hoang *et al.*, 1998) |
| pMMB67EH-Gm | Broad-host-range bacterial cloning vector, *tac* promoter (GmR) | This study |
| pMMB67-RocS1 | pMMB67EH-GW harbouring the *rocS1* gene from Gateway library (GmR) | Kulasekara et al., 2005) |
| pMMB67EH | Broad host range bacterial cloning vector, *tac* promoter (ApR/CbR) | (Sivaneson *et al.*, 2011) |
| pMMB67-RocS2 | pMMB67HE harbouring the *rocS2* gene from Gateway library (ApR/CbR) | (Sivaneson *et al.*, 2011) |
| pEXG2 | Allelic exchange vector (GmR ) | (Rietsch *et al.*, 2005) |
| pEXG2 Δ*PA4080* | pEXG2 carrying SLIC fragment for *rocA3* deletion (GmR) | This study |
| pEXG2 PAO1Δ*rocA1* | pEXG2 carrying SLIC fragment for *rocA1* deletion (GmR) | This study |
| pEXG2 PAO1Δ*rocA2* | pEXG2 carrying SLIC fragment for *rocA2* deletion (GmR) | This study |
| pEXG2 *PA4080-D58A* | pEXG2 carrying SLIC fragment for *rocA3* mutation (GmR) | This study |
| pJN105 | P*araBAD* (pBAD) transcriptional fusion vector (GmR) | (Newman and Fuqua, 1999) |
| pJN105-PA4080 | pJN105 carrying P*BAD-rocA3* transcriptional fusion (GmR) | This study |
| pJN105-rocR | pJN105 carrying P*BAD-rocR* transcriptional fusion (GmR) | This study |
| pJN105-rocR3 | pJN105 carrying PBAD-(IHMA87) rocR3 transcriptional fusion (GmR) | This study |
| miniCTX-TrrnB-lacZ | Site-specific integrative plasmid with promoter less-lacZ and strong rrnB terminator (attP site, FRT, TcR) | Elsen et al. (2023) |
| pCTXter-PcupB1-lacZ | miniCTX-TrrnB-lacZ harboring the cupB1 promoter fused to lacZ (attP site, FRT, TcR) | This study |
| pCTXter-PcupC1-lacZ | miniCTX-TrrnB-lacZ harboring the cupC1 promoter fused to lacZ (attP site, FRT, TcR) | This study |
| pCTXter-PPA4080-lacZ | miniCTX-TrrnB-lacZ harboring the rocA3 promoter fused to lacZ (attP site, FRT, TcR) | This study |
| pCTXter-PmexA-lacZ | miniCTX-TrrnB-lacZ harboring the mexA promoter fused to lacZ (attP site, FRT, TcR) | This study |
| pCTXter-PlepB-lacZ | miniCTX-TrrnB-lacZ harboring the lepB promoter fused to lacZ (attP site, FRT, TcR) | This study |
| pKT15 | Cloning and expression vector, encodes the T25 fragment (amino acids 1–224 of CyaA) (KmR) | (Karimova et al., 1998) |
| pKT25-HptA | Fusion of hptA to cya gene T25 fragment in pKT25 (KmR) | This study |
| pKT25-HptB | Fusion of hptB to cya gene T25 fragment in pKT25 (KmR) | This study |
| pUT18c | Cloning and expression vector, encodes the T18 fragment (amino acids 225–399 of CyaA) (ApR) | (Karimova et al., 1998) |
| pUT18c-2583-D1 | Fusion of sequence encoding D1 domain (834-992 aa) of PA2583 to cya gene T18 fragment in pUT18C (ApR) | This study |
| pUT18c-SagS-D1 | Fusion of sequence encoding D1 domain (663-786 aa) of SagS to cya gene T18 fragment in pUT18C (ApR) | This study |
| pUT18c-RocA1-D2 | Fusion of sequence encoding D2 domain of RocA1 to cya gene T18 fragment in pUT18C (ApR) | (Kulasekara et al., 2004) |
| pUT18c-RocA2-D2 | Fusion of sequence encoding D2 domain of RocA2 to cya gene T18 fragment in pUT18C (ApR) | (Sivaneson et al., 2011) |
| pUT18c-RocA3-D2 | Fusion of sequence encoding D2 domain (1-134 aa) of RocA3 to cya gene T18 fragment in pUT18C (ApR) | This study |
| pUT18c-RocR-D2 | Fusion of sequence encoding D2 domain of RocR to cya gene T18 fragment in pUT18C (ApR) | (Kulasekara et al., 2004) |
| pUT18c-RocR3-D2 | Fusion of sequence encoding D2 domain (1-136 aa) of RocR3 to cya gene T18 fragment in pUT18C (ApR) | This study |

**Table S2. Primers used in this work**

| **Name** | **Sequence (5’ => 3’)** | **Use** |
| --- | --- | --- |
| **pEXG2-mut-PA4080-sF1** | GGTCGACTCTAGAGGATCCCC TCTGAGGGATATGCAAGTTTCCT | *rocA3* deletion |
| **pEXG2-mut-PA4080-sR1** | GCGATGTAGGGACATCCTTGC | *rocA3* deletion |
| **pEXG2-mut-PA4080-sF2** | GCAAGGATGTCCCTACATCGC TGCCGGATGCCCTGACGGAT | *rocA3* deletion |
| **pEXG2-mut-PA4080-sR2** | ACCGAATTCGAGCTCGAGCCC GATGTGGCGGAAATCGACGCC | *rocA3* deletion |
| **pEXG2-mut-rocA1-sF1** | GGTCGACTCTAGAGGATCCCC GATGTCGACGTGTCCGCAGG | *rocA1* deletion |
| **pEXG2-mut-rocA1-sR1** | CAGGACGGTATGCATAAATTCGA | *rocA1* deletion |
| **pEXG2-mut-rocA1-sF2** | GAATTTATGCATACCGTCCTG TCGCTGATCCACTGACCTGGC | *rocA1* deletion |
| **pEXG2-mut-rocA1-sR2** | ACCGAATTCGAGCTCGAGCCC GAATTTCCGGCCGACCGAGG | *rocA1* deletion |
| **pEXG2-mut-rocA2-sF1** | GGTCGACTCTAGAGGATCCCC GAAGCTCTAAGCGATAGCGCC | *rocA2* deletion |
| **pEXG2-mut-rocA2-sR1** | TATCAGGATACGGCTCATTGTCT | *rocA2* deletion |
| **pEXG2-mut-rocA2-sF2** | ACAATGAGCCGTATCCTGATA GAGTTCGCCAAACGCAATACGC | *rocA2* deletion |
| **pEXG2-mut-rocA2-sR2** | ACCGAATTCGAGCTCGAGCCC GTTGCAGGCCAGCAGGCGAC | *rocA2* deletion |
| **pEXG2-mut-PA4080-sF1** | GGTCGACTCTAGAGGATCCCC TCTGAGGGATATGCAAGTTTCCT | *rocA3-D58A* mutation |
| **pEXG2-PA4080-D58A-sR1** | GGCATCGTATAGGCGACGATAAC | *rocA3-D58A* mutation |
| **pEXG2-PA4080-D58A-sF2** | GTTATCGTCGCCTATACGATGCC | *rocA3-D58A* mutation |
| **pEXG2-mut-PA4080-sR2** | ACCGAATTCGAGCTCGAGCCC GATGTGGCGGAAATCGACGCC | *rocA3-D58A* mutation |
| **pCTXter-cupB1-lacZ-sF** | GATATCGAATTCCTGCAGCCC TCCATCCGGAATGCGAGTGGG | *PcupB1-lacZ fusion* |
| **pCTXter-cupB1-lacZ-sR** | GCTAGTTAGTTAGGATCCCCC CATCTGATTTCCTTTGGAGTTGTG | *PcupB1-lacZ fusion* |
| **pCTXter-cupC1-lacZ-sF** | GATATCGAATTCCTGCAGCCC AGGCAAACTAAGTGCGTTCGAAA | *PcupC1-lacZ fusion* |
| **pCTXter-cupC1-lacZ-sR** | GCTAGTTAGTTAGGATCCCCC CATGATTGAGCTTCCTTTTGACAG | *PcupC1-lacZ fusion* |
| **pCTXter-PA4080-lacZ-sF** | GATATCGAATTCCTGCAGCCC GCTCGTCCTGGGTCTCGTG | *ProcA3-lacZ fusion* |
| **pCTXter-PA4080-lacZ-sR** | GCTAGTTAGTTAGGATCCCCC ATCCTTGCGTCTCCTTAAGC | *ProcA3-lacZ fusion* |
| **pCTXter-mexA-lacZ-sF** | GATATCGAATTCCTGCAGCCC AGCTCGCGGATCTTCCG | *PmexA-lacZ fusion* |
| **pCTXter-mexA-lacZ-sR** | GCTAGTTAGTTAGGATCCCCC ATAGCGTTGTCCTCATGAGC | *PmexA-lacZ fusion* |
| **pCTXter-lepB-lacZ-sF** | GATATCGAATTCCTGCAGCCC TCTAGCGCTGCGTCCTGCC | *PlepB-lacZ fusion* |
| **pCTXter-lepB-lacZ-sR** | GCTAGTTAGTTAGGATCCCCC ATGAGGCCCGTACGGAGC | *PlepB-lacZ fusion* |
| **pJN105-PA4080-sF** | CTAGCGAATTCCTGCAGCCC TTTCCTGGTAACGCTTTCGCCT | *rocA3 expression* |
| **pJN105-PA4080-sR** | CTAGAACTAGTGGATCCCCC TCAGGGCATCCGGCAATAGCC | *rocA3 expression* |
| **pJN105-rocR-sF** | CTAGCGAATTCCTGCAGCCC AACACAGCAGTATATCGCCGGT | *rocR expression* |
| **pJN105-rocR-sR** | CTAGAACTAGTGGATCCCCC TAGAGGCAGCCGAGGCCAG | *rocR expression* |
| **pJN105-rocR3-sF** | CTAGCGAATTCCTGCAGCCC ACAATCGCTCCACCATAAAGATTA | *IHMA87 rocR3 expression* |
| **pJN105-rocR3-sR** | CTAGAACTAGTGGATCCCCC GTTGATATCGCAACGACCGCC | *IHMA87 rocR3 expression* |
| **pKT25-hptA-sR** | TACTTAGGTACCCGGGGATC TCTAGAGCTTTGCCAATTCGGTT | *pKT25 cloning for BTH* |
| **pKT25-hptA-sF** | CTGCAGGGTCGACTCTAGAG ATGAAAGAGCTTGGTTCGGAATC | *pKT25 cloning for BTH* |
| **pKT25-hptB-sR** | TACTTAGGTACCCGGGGATC TCAGCGATAGCGCTGACGTTC | *pKT25 cloning for BTH* |
| **pKT25-hptB-sF** | CTGCAGGGTCGACTCTAGAG ATGTCCGCGCCGCATCTCGA | *pKT25 cloning for BTH* |
| **pUT18c-PA2583 D1-sR** | GAGCTCGGTACCCGGGGATC TCATTGCGAATCGACGCTCGT | *pUT18c cloning for BTH* |
| **pUT18c-PA2583 D1-sF** | ACTGCAGGTCGACTCTAGAG ACCGAGCCTCCGCCCAGC | *pUT18c cloning for BTH* |
| **pUT18c-sagSD1-sF** | ACTGCAGGTCGACTCTAGAG ACCCGGGTCCTGCTGGTGGA | *pUT18c cloning for BTH* |
| **pUT18c-sagSD1-sR** | GAGCTCGGTACCCGGGGATC CTAGTCGCTCGCGGTGAGCG | *pUT18c cloning for BTH* |
| **pUT18c-rocA2D2-sF** | ACTGCAGGTCGACTCTAGAG ATGAGCCGTATCCTGATAGTCG | *pUT18c cloning for BTH* |
| **pUT18c-rocA2D2-sR** | GAGCTCGGTACCCGGGGATC CTACGAGCTGGGAAAGTAACTGTAT | *pUT18c cloning for BTH* |
| **pUT18c-PA4080D2-sF** | ACTGCAGGTCGACTCTAGAG ATGTCCCTACATCGCATTCGC | *pUT18c cloning for BTH* |
| **pUT18c-PA4080D2-sR** | GAGCTCGGTACCCGGGGATC TTACGGCGAGAAATACTGGTTTCG | *pUT18c cloning for BTH* |
| **pUT18c-rocR3D2-sF** | ACTGCAGGTCGACTCTAGAG ATGCGGCAGATCAGCGTCCT | *pUT18c cloning for BTH* |
| **pUT18c-rocR3D2-sR** | GAGCTCGGTACCCGGGGATC CTAAAGAGCGGGCGGAAGCAGC | *pUT18c cloning for BTH* |
